# Investigating interspecific quorum sensing influence on cocoa fermentation quality through defined microbial cocktails

**DOI:** 10.1101/2022.06.14.496151

**Authors:** O.G.G. Almeida, M. G. Pereira, R. L. Bighetti-Trevisan, E.S. Santos, E. G. De Campos, G.E. Felis, L.H.S. Guimarães, M.L.T.M Polizeli, B. S. De Martinis, E.C.P. De Martinis

**Affiliations:** Universidade de São Paulo, Faculdade de Ciências Farmacêuticas de Ribeirão Preto. Departamento de Análises Clínicas, Toxicológicas e Bromatológicas; Universidade do Estado de Minas Gerais, Unidade Passos; Universidade de São Paulo, Faculdade de Odontologia de Ribeirão Preto. Departamento de Biologia Básica e Oral.; Appalachian State University, Department of Chemistry and Fermentation Sciences, Boone, NC, United States; University of Verona, Department of Biotechnology, Verona, Italy; Universidade de São Paulo, Faculdade de Filosofia Ciências e Letras de Ribeirão Preto. Departamento de Biologia.; Universidade de São Paulo, Faculdade de Filosofia Ciências e Letras de Ribeirão Preto. Departamento de Química.

**Keywords:** cocoa fermentation, quorum sensing, starter cultures, *luxS* gene

## Abstract

The fermentation of cocoa beans is a key process to supply high quality ingredients for the chocolate industry. In spite of several attempts to obtain standardised microbial cultures for cocoa fermentation, it is still a spontaneous process. It has been suggested lactobacilli present potential for quorum sensing (QS) regulation in cocoa fermentation, and in the present research, laboratory scale fermentations were carried out to further elucidate possible QS influence on microbial shifts and fermented seeds quality. The experimental design comprised the 96 hours-fermentations designated as F0 (control), F1 (yeasts, lactic acid bacteria, and acetic acid bacteria), F2 (yeasts and acetic acid bacteria), F3 (yeasts only), with evaluation of the microbial succession by plate counting, determination of enzymatic activities by classical methods and qualitative evaluation of flavour compounds by gas-chromatography (GC-MS) with headspace sampling. Besides, QS was estimated by quantification of the expression of *luxS* genes by Reverse Transcriptase Real Time PCR analysis using selected primers. The results demonstrated that microbial successions were displayed in lab conditions, but no statistical difference in terms of microbial enumeration and α-diversity metrics were observed among the experimental and control fermentations. Moreover, enzymatic activities were not correlated to the total microbiota, indicating the seeds’ endogenous hydrolases protagonist enzymes secretion and activity. Regarding *luxS* genes measuring for the species *Lactiplantibacillus plantarum* and *Limosilactobacillus fermentum*, genes were active in fermentation in the start to the end phase and to the beginning to the middle phase of fermentation, respectively. Correlation analysis among *luxS* expression and volatile metabolites evidenced *Lp. plantarum* association with detrimental compounds for fermentation quality. This data contributes to our previous research which monitored fermentations to survey enzymatic changes and QS potential along the process and sheds light of QS-related strategies of lactobacilli dominance in cocoa fermentations.

## Introduction

Cocoa fermentation is a spontaneous process characterised by a succession of yeasts, lactic acid bacteria (LAB) and acetic acid bacteria (AAB), which are responsible for generating desirable sensory characteristics in the raw material for chocolate production, encompassing colour, aroma, flavour, and texture (De Vuyst & Weckx, 2016). However, such biological process is complex, as the microbial composition may vary geographically and according to the fermentation methods and fermentative processes used (Bortolini et al., 2016). Although many studies indicated *Lactiplantibacillus plantarum* and *Limosilactobacilus fermentum* species dominate in several fermentations carried out in diverse geographic regions, the determination of a core microbiome for cocoa fermentation is challenging (Viesser et al., 2021). In Brazil, for instance, it was observed a superior diversity of LAB and AAB in comparison to Ghana which exhibited a varied repertoire of LAB instead. On the other hand, other producing regions such Nicaragua, Colombia, Cameroon, and Ivory Coast showed lower bacterial diversities due to the paucity of NGS data (Viesser et al., 2021).

Besides, little is still known about the intraspecific variation among the strains of cocoa dominant species for the selection of those with metabolic repertoires of interest in order to standardise the process (Ouattara & Niamké, 2021). The search for a starter culture is not a novelty, as so many works have proposed a multitude of candidate strains with interesting metabolic traits (Farrera et al., 2021; Magalhães da Veiga Moreira et al., 2017; Ooi et al., 2020; Saunshi et al., 2020; Visintin et al., 2017). However, the criteria for the selection of these microorganisms are yet a subject of much discussion and there is no guideline for this, since they are based, mostly, on the correlation of the sensory profiles generated after the inoculation of strains in fermentations, inhibition of detrimental and pathogenic microbiota and mobilisation of pulp components (Chagas Junior et al., 2021).

Among the characteristics of a well-defined starter culture is its potential to tolerate adverse conditions and to compete with undesirable microorganisms, replacing them and assuring safety. Microorganisms must overcome stressors by the expression of heat shock proteins, production of exopolysaccharides, releasing of antimicrobial compounds and harbouring specific metabolic pathways that assure their permanency in an environment (De Roos & De Vuyst, 2018; García-Ríos et al., 2021; Ogunremi et al., 2022; Romanens et al., 2019). The robustness of a microbial species resides in all these characteristics gathered in the fermentation.

Recent studies relate quorum sensing (QS) to the adaptation of bacteria to stressful conditions. QS is a phenomenon associated to the synchronisation of gene expression in bacteria in response to the density of bacterial populations leading to a multitude of downstream metabolic responses such as phenotypic modulation and physiological changes to maintain homeostasis (Johansen & Jespersen, 2017; Wu et al., 2020; Yang et al., 2021). The *luxS* gene is the universal gene marker for interspecific quorum sensing as it is responsible for the conversion of 4,5-dihydroxy-2,3-pentanedione (DPD) molecule to the autoinducer 2 (AI-2) that will sensitise adjacent cells in the environment, tiggering QS. The AI-2 is produced in the methyl-cycle from the S-adenosylmethionine (SAM) precursor molecule. The pathway starts by the transferring of a methyl group from SAM to methyl-transferases and substrates producing S-adenosylhomocysteine (SAH). Then, SAH has an adenine removed by a nucleosidase (pfs) enzyme leading to the production of a S-ribosyl homocysteine (SRH), whose reaction is catalysed by the LuxS enzyme. The SRH is finally converted into homocysteine and DPD. As this last is an unstable molecule, it can spontaneously cyclize into several DPD derivatives and sensibilize several bacteria (Wu et al., 2020; Yang et al., 2021). As this gene is widespread in the bacterial kingdom as it is part of the activated methyl-cycle, it could be a clue of a universal interspecies communicator (Tobias et al., 2020), being and excellent marker to assess bacterial communication by *in silico* methods.

In this sense, a metagenomic study of spontaneous cocoa fermentation carried out by our research group showed that lactic acid bacteria, with emphasis on lactobacilli, were correlated to higher amounts of *luxS*-derivate gene reads along cocoa fermentation (Almeida et al., 2020). That study showed that as the lactobacilli load increased, the *luxS* gene counts were also increased. Another study, carried out on kimchi’s fermentation, showed that the most dominant LAB bacteria showed the expression of *luxS*, while non-dominant bacteria did not (Park et al., 2016). Other studies also demonstrate that the expression of the *luxS* gene in LAB can result in adaptive benefits in the face of unfavourable conditions related to low pH, high temperatures and nutrient depletion (Gu et al., 2018; Jiang et al., 2021).

Thus, in view of starter cultures drawing QS characterization of strains, accompanied with sensorial screening and other metrics already used in literature, could be a valuable tool to select robust candidates in the aim of maintaining products identity balanced with standardisation. On the other hand, the information relating QS to the quality of fermented foods is scarce and must be evaluated in depth. In this context, this work aims to determine through lab-scale fermentation the influence of interspecific QS on the dominance of LAB in cocoa fermentation and if it influences the quality of fermented seeds by the use of defined yeasts, LAB, and AAB cocktails. To test this hypothesis, production of volatile metabolites, enzymatic activity changes, and microbial composition were evaluated by chemistry based and NGS methods. Besides, *luxS* gene expression was monitored, quantified along the fermentations, and analysed by correlation and regression to answer the following question: “Does quorum sensing play a role in microbial shifts along spontaneous fermentation of cocoa beans?”.

## Materials and methods

### Selected strains

All the strains used in this study were obtained from a spontaneous cocoa fermentation carried out in Bahia state, Brazil (Almeida et al. 2020). The fungal strains selected were *Pichia kudriavzevii* strain PCA1, *Pichia kluyveri* strain PCA4 and four *Saccharomyces cerevisiae* with distinct genotypes as determined by the microsatellite amplification (Vaudano & Garcia-Moruno, 2008), strains F3, F6, F11 and F12. These strains were selected since *Pichia* and *Saccharomyces* species are often reported in cocoa fermentation.

Regarding LAB, the strains *Lactiplantibacillus plantarum* Lb2 and *Limosilactobacillus fermentum* Lb1 were selected as they were characterised in terms of potential interspecific QS by genomic-centred analysis (Almeida et al., 2021). Finally, five genotypically diverse *Acetobacter senegalensis* strains named MRS7, GYC10, GYC12, GYC19 and GYC27 were selected due to their genetic potential to adapt in harsh fermentative environments (Almeida et al., 2022).

### Inoculum preparation

The yeasts were cultured on a 50 ml of yeast extract (1%), peptone (2%) and dextrose (2%) medium (YPD) and incubated for 24h at 30°C under shaker agitation at 110 rpm. LAB were cultured on a 50 ml of De Man, Rogosa & Sharpe (MRS, Oxoid, Basingstoke, UK) broth, incubated at 30°C for three days in anaerobic jars. AAB were cultured on a 50 ml of a modified broth of glucose (5%) and yeast extract (1%) (GYC medium) and incubated for 24h at 30°C under shaker agitation at 110 rpm. Then, the strains were enumerated to standardise the inoculum load on fermentations. A concentration of 1×10^6^ UFC/g was determined using a growth curve as the optimal concentration for each yeast and bacterial species selected. The strains were recovered from their culture media by centrifugation for 15 minutes at 7,500g. Then, each strain’s pellet was resuspended in saline solution and centrifuged for 15 minutes at 7,500g twice. Dedicated volumes were added in sterile Boeco® flasks to compose the cocktails. The yeast, LAB and AAB cocktails totalized a volume of 300 ml, 200 ml and 100ml, respectively.

### Fermentation experimental design

Cocoa fruits (mix of cultivars Criolo and Forastero) were obtained from “Companhia de Entrepostos e Armazéns Gerais de São Paulo - CEAGESP” distributor located at Ribeirão Preto city (Brazil) and were used in all fermentations. To carry out the fermentation, the fruits were opened with a sterile stainless-steel knife to perform the peeling. A total of 5 kg of fresh-harvested cocoa seeds was placed in each box measuring 40 cm long, 12 cm high and 29 cm width which were partially covered with the lid along the entire fermentation. This was considered the start time of fermentation (time 0h). After the first 48 h, the seeds were tuned for aeration, being this process repeated every 24h as performed in field conditions.

The experimental design was composed by a control fermentation, named F0, which was composed only by cocoa seeds without replicates, and three main fermentations performed in biological duplicates, named: F1, F2 and F3. The experimental design was depicted in Fig. 1. The C1 cocktail was prepared in a volume of 300 ml of *Pichia kudriavzevii* PCA1, *Pichia kluyveri* PCA4, and *Saccharomyces cerevisiae* strains F3, F6, F11, and F12 in equimolar concentrations of 1×10^6^ yeasts/g of each species. The C2 cocktail was prepared in a volume of 200 ml of *Lactiplantibacillus plantarum* Lb2 and *Limosilactobacillus fermentum* Lb1 in equimolar concentrations of 1×10^6^ bacteria/g of seeds of each species. The C3 cocktail was prepared in a volume of 100 ml of *Acetobacter senegalensis* strains MRS7, GYC10, GYC12, GYC19 and GYC27 in concentrations of 1×10^6^ bacteria/g of each species.

**Fig. 1.**
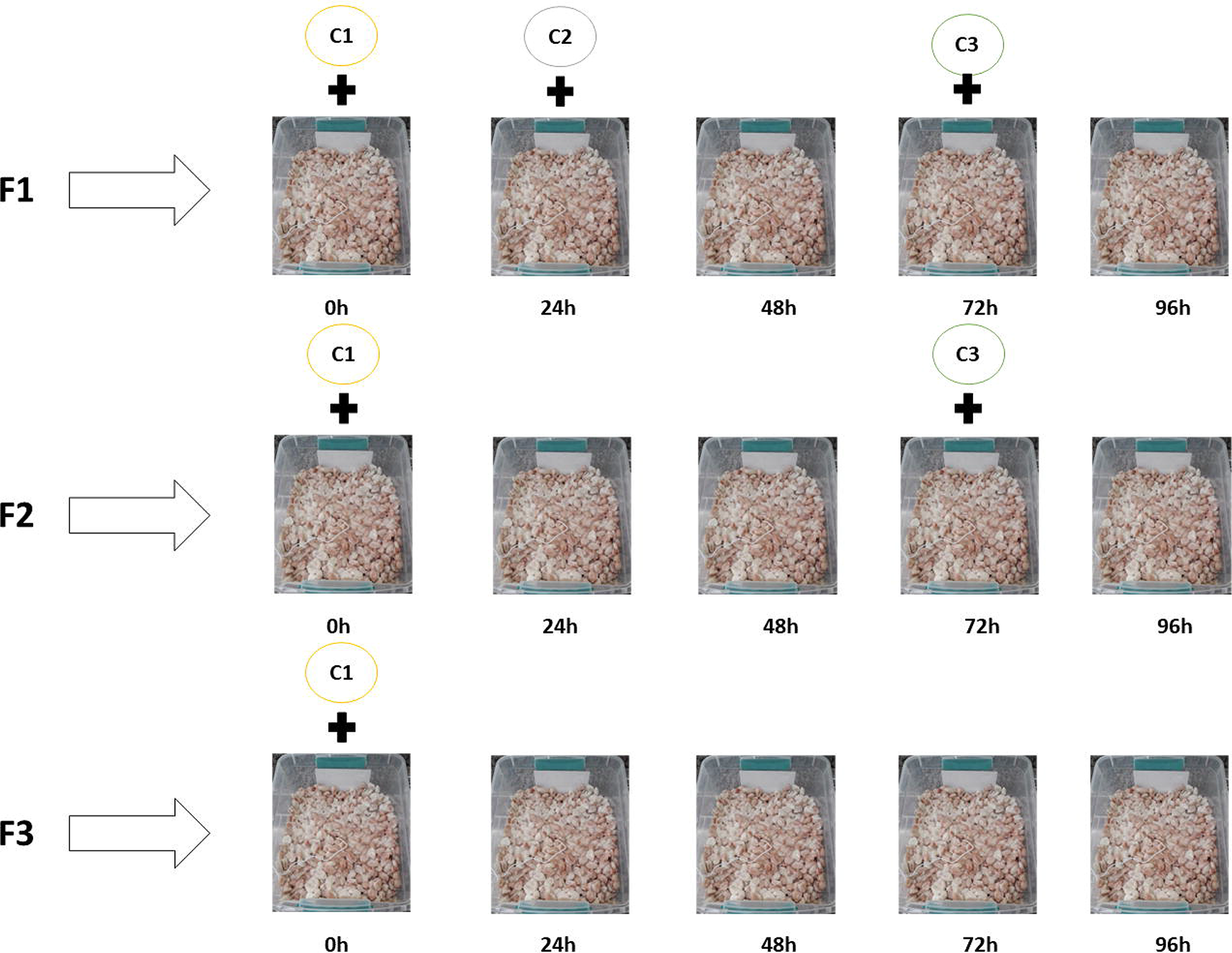
Experimental design. Schema of inoculation of the defined cocktails C1 (*Pichia kudriavzevii* PCA1, *Pichia kluyveri* PCA4, *Saccharomyces cerevisiae* strains F3, F6, F11, and F12), C2 (*Lactiplantibacillus plantarum* Lb2 and *Lm. fermentum* Lb1), and C3 (*Acetobacter senegalensis* strains MRS7, GYC10, GYC12, GYC19 and GYC27) in F1, F2 and F3 fermentations.

In the F1 fermentation, all cocktails C1, C2, and C3 were added at the times 0h, 24h, and 72h, respectively. In the fermentation F2, the C2 cocktail was not inoculated, but only the C1 and C3 cocktails were added at the times 0h and 72h, respectively. Finally, F3 fermentation was inoculated only with the C1 cocktail at the time 0h. Boxes temperature and pH were measured continuously using a digital thermohygrometer (Kasvi, Paraná, Brazil) and a pHmeter (Quimis, São Paulo, Brazil).

### Sampling and enumeration of microorganisms from lab-scale cocoa fermentations

During the fermentations the samples were collected aseptically in intervals of 0h, 24h, 48h, 72h and 96h for enumeration of presumptive yeasts, LAB, and AAB. For this, selective media for these microorganisms were used: Amphotericin B (Sigma-Aldrich, Massachusetts, USA) (5mg/l) in MRS (Oxoid) and GYC culture media and chloramphenicol (Sigma-Aldrich) (200 mg/l) in YPD. 225 ml of 0.1% peptone water was added to 25 g of each sample and then it was manually homogenised for 2 min. Then, serial dilutions were performed in 0.1% peptone water. After the process, the samples were plated in three different types of culture media (GYC, MRS (Oxoid) and YPD), containing antibiotic and/or antifungal. The plates were incubated for 48 h and then counting was performed.

### Crude extract preparation and enzymatic activities quantification

To determine the total protein content and enzymatic activity catalysed by cocoa microbiota, 17 g of seeds were homogenised in 30 ml of distilled water by vortex for complete homogenization of the pulp. This mixture was placed in 50 ml Falcon® tubes which were kept refrigerated at 4°C up to 8°C. Enzyme activities were determined using the direct substrates for the enzyme. One unit (U) was defined as the amount of enzyme necessary to hydrolyze 1 μmol of substrate per minute under the previously described conditions.

Lipase activity was determined using *p*-nitrophenylpalmitate (Sigma-Aldrich) as substrate (Pencreac’h & Baratti, 1996). Pectinase activity by the 3’,5’-dinitrosalicylic acid (DNS) (Sigma-Aldrich) method (Miller, 1959), using 1% galacturonic acid solution dissolved in 50 mM sodium acetate buffer, pH 5.0. Amylase activity was determined using the reaction by the DNS method (Miller, 1959), using 1% starch solution dissolved in 50 mM sodium acetate buffer, pH 5.0. Arabinase activity was dosed using an unbranched arabinan as substrate through the formation of reducing sugars by the DNS method (Miller, 1959). Arabinofuranosidase activity was detected using the batch method (Kersters-Hilderson et al., 1982), using the synthetic substrate and *p*-nitrophenyl-α-L-arabinofuranoside (Sigma-Aldrich) (PNP-ara). Cellulase activity was performed with the substrate avicel 1% (w/v) through the formation of reducing sugars by the DNS method (Miller, 1959). Invertase activity was measured using sucrose as substrate 1% (w/v) through the formation of reducing sugars by the DNS method (Miller, 1959). β-glucanase activity was measured using β-glucan syrup 0.5% (w/v) as substrate through the formation of reducing sugars by the DNS method (Miller, 1959). Endoglucanase activity was measured using Carboximetil celulose (Sigma-Aldrich) (CMC) 1% (w/v) as substrate through the formation of reducing sugars by the DNS method (Miller, 1959). Xyloglucanase activity was measured using xyloglucan 1% (w/v) as substrate through the formation of reducing sugars by the DNS method (Miller, 1959). Mannanase activity was measured using “Locust bean gum” 1% (w/v) as substrate through the formation of reducing sugars by the DNS method (Miller, 1959). Xylanase activity was measured using xylan Beechwood (Sigma-Aldrich) 1% (w/v) as substrate through the formation of reducing sugars by the DNS method (Miller, 1959). Esterase activity was measured using *p*-nitrophenyl-acetate (PNP-acetate) (Sigma-Aldrich) 1% (w/v) as substrate through the formation of reducing sugars by the DNS method (Miller, 1959). Cellobiohydrolase activity was measured using the synthetic substrate *p*-nitrophenyl-cellobioside (Sigma-Aldrich) (PNP-cellobioside). β-glucosidase activity was measured using the synthetic substrate *p*-nitrophenyl-glucopyranoside (Sigma-Aldrich) (PNP-Glu). β-xylosidase activity was measured using the synthetic substrate *p*-nitrophenyl-xylopyranoside (Sigma-Aldrich) (PNP-xylo).

Protein amounts were determined using the Bradford (1976) method and a bovine serum albumin curve as a standard reference.

### Volatile compounds identification

Prior to analysis, frozen cocoa samples (pulp and seed) were removed from the freezer, weighted (1.00 to 1.05 g) and inserted into a headspace vial. Samples were manually subjected to incubation in a dry block heater for 30 minutes, at 90°C. After the incubation, 0.5 mL of the vapour phase was collected with a gas tight syringe and manually injected into the GC-MS for analysis.

Analyses were performed using an Agilent (Santa Clara, CA, United States) 7890A GC coupled to an Agilent 5975C MS (Santa Clara, CA, United States). A capillary column HP-5MS (30 m X 0.25 mm, 0.25 mm) was used for the chromatographic separation. The method used in these analyses was based on previously published methods available in the literature (Instituto Adolfo Lutz, 2008; Tait et al., 2014). The separation was performed using a temperature program as follows: 40°C for 5 min, increased to 220°C at 8°C/min e isothermal for 2 min. Injection was performed at 250°C, in split mode, with a split ratio of 1:10. Helium was used as carrier gas, at a flow rate of 1 ml/min. The mass spectrometer operated in Full Scan mode (m/z 50 – 650). The temperatures of the MS source and quadrupole were 230 and 150°C, respectively.

Data obtained from the analyses were performed using the AMDIS from NIST (Version 2.66, August 2008), available within the GC-MS software. The following settings were adopted for data analysis and processing using the AMDIS software: (a) type of analysis: simple; (b) minimum match factor set at 60 and (c) parameters of resolution, sensitivity and shape requirements were used as the default setting (Medium). Library used for the searches was the NIST Mass Spectral Library (Version 2.0, October 2009). The criteria for the search in the NIST library was a match of 700 or higher (NIST, 2008). Only metabolites detected in both replicates were summarised and reported.

### Metagenomic DNA extraction and sequencing

Metagenomic DNA extraction was performed according to the protocol described by our research group in the reference (Almeida et al., 2020). In summary, three random seeds were selected, and their pulps were scraped. Then, DNA extraction was performed according to the recommendations of the manufacturer of the Zymobiomics DNA MINI KIT (California, USA). The extracted DNA was quantified by Qubit fluorometer (ThermoFisher) and nanodrop (ThermoFisher) and sent to the “BPI Biotecnologia EPP’’ facility for 16S *rRNA* V3-V4 region (forward primer: 5’ TCGTCGGCAGCGTCAGATGTGTATAAGAGACAGCCTACGGGNGGCWGCAG3’; reverse primer: 5’GTCTCGTGGGCTCGGAGATGTGTATAAGAGACAGGACTACHVGGGTATCTAAT CC3’) and *ITS* region amplification (primer 86F: 5’ TCGTCGGCAGCGTCAGATGTGTATAAGAGACAGGTGAATCATCGAATCTTTGAA3’; 4R: 5’ GTCTCGTGGGCTCGGAGATGTGTATAAGAGACAGTCCTCCGCTTATTGATATGC3’) and sequencing on the MiniSeq platform (Illumina®) to generate paired-end (PE) 2×250 bp reads following the facility’s protocols. Raw sequencing data was made publicly available on NCBI Bioprojects PRJNA842267 and PRJNA842340.

### Total RNA extraction for qRT-PCR

Total RNA extraction was performed according to the protocol described by Verce et al. (2021), with modifications. Briefly, three cocoa beans were homogenised in 30 ml of RNAprotect (Qiagen, Venio, The Netherlands) in a 50 ml falcon and vortexed for 30 seconds to disrupt the mucilaginous pulp from the seeds. Then, the seeds were removed with the aid of sterile forceps. Then, the tubes were centrifuged at 6,000g for 30 minutes. Subsequently, the supernatant was discarded, and 2 ml of sorbitol wash buffer (1.5 M sorbitol, 50 mM Tris-base and nuclease-free water) were added, followed by vigorous vortex, and centrifugation for 10 minutes at 6,000g. The supernatant was discarded and 3.5 ml of RLT buffer with B-mercaptoethanol (10 µl for every 350 µl of RLT buffer – Qiagen) was added. The tubes were vortexed for 20 seconds and then centrifuged for three minutes at maximum speed (about 10,000g). Then, the supernatant was eluted on the RNeasy Mini Kit (Qiagen) gDNA strip elution column until the entire contents of each tube were exhausted. Then, 2.0 ml of 70% alcohol was added to the eluted volume, which was gently homogenised with the pipette. Finally, the remaining steps were performed according to the RNeasy Mini Kit manufacturer’s instructions for RNA obtaining. The integrity of the extracted RNA was evaluated in a 1% (m/v) agarose gel and the material was quantified using the nanodrop (ThermoFisher, Massachusetts, USA).

### *luxS* gene quantification by qRT-PCR

For relative quantification by qRT-PCR, the *luxS* genes of *Lm*. *fermentum* and *Lp. plantarum* species were amplified in parallel with the respective 16S *rRNA* genes for gene expression normalisation. The chosen primers were already described in literature (Almeida et al., 2021; Gu et al., 2018; Schwendimann et al., 2015) and are presented in the Table 1. The cDNA preparation was performed following the instructions of the manufacturer of the ProtoScript II Reverse Transcriptase (New England Biolabs, Massachusetts, USA). Random primers (ThermoFisher) were used to initiate the reverse transcription reaction. Quantification of relative *luxS* gene expression was performed by real-time qPCR using the qPCRBIO SyGreen Mix (PCR Biosystems, Pennsylvania, USA), according to the manufacturer’s instructions. The reactions were carried out in triplicate in the Realplex^4^ Epgradient Mastercycler (Eppendorf, Hamburg, Germany). The 2^-ΔΔCt^ method (Livak & Schmittgen, 2001) was applied to calculate the relative expression levels of *luxS* genes, using the 16S *rRNA* Ct values as normalizer.

**Table 1.**
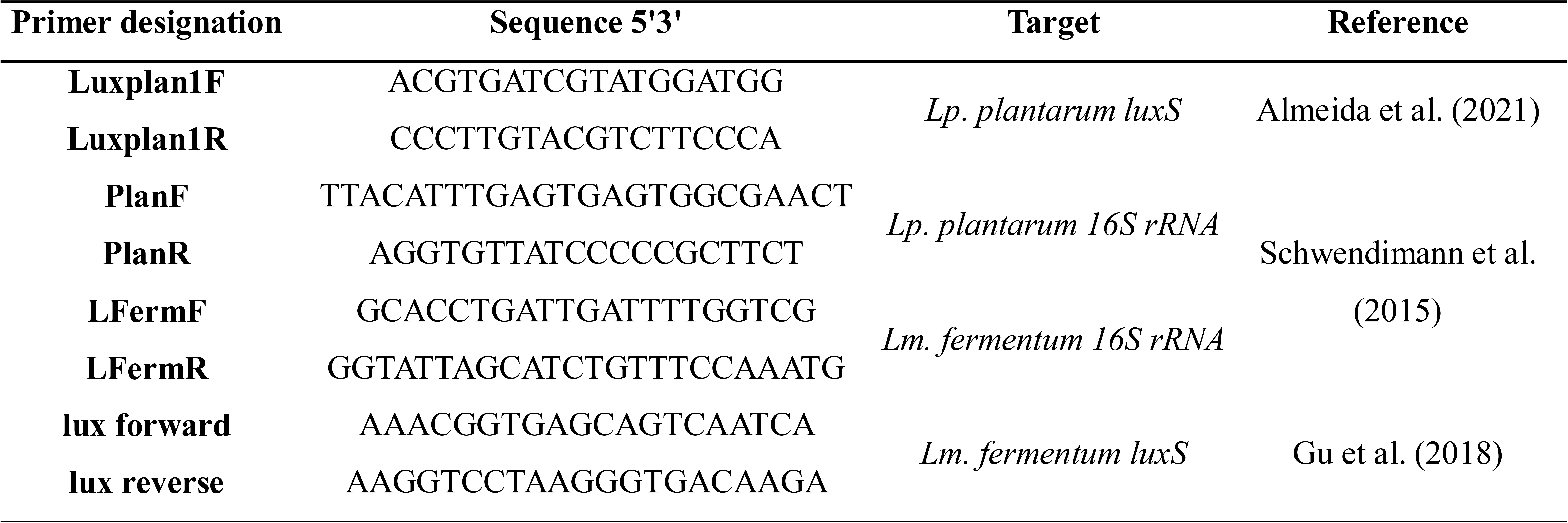
Primers sequences for qRT-PCR reactions.

### Bioinformatic analysis of microbial composition

The high-quality forward reads (R1) were selected for downstream analysis on Qiime2 environment (Bolyen et al., 2019). Initially, the delivered raw reads were assessed in terms of quality using the bbduk tools (Bushnell, 2014) to remove adapters/barcodes fragments and reads with phred scores lower than 30 (parameters: hdist=1 tpe tbo qtrim=rl trimq=30 maq=30). Then, the reads were imported and processed with the dada2 (Callahan et al., 2016) plugin for chimera removal and de-replication. The taxonomy was assigned by training the full 16S *rRNA* region of the 99% SSU NR database obtained from the SILVA repository (Quast et al., 2013). The same initial steps were followed for fungi taxonomic assignment, but the database chosen was the UNITE 99% NR version 10.05.2021 (Nilsson et al., 2019).

The processed ASVs (amplicon sequence variants) data was imported to the RStudio environment using the R package qiime2R (Jordan & Bisanz, 2018). Data visualisation (i.e., alpha-diversity, beta-diversity, and composition) was generated using dedicated microbiome packages such as phyloseq (McMurdie & Holmes, 2013), qiime2R (Jordan & Bisanz, 2018), and microbiomeutilities (Shetty & Lahti, 2020). Statistical analysis was performed using the Wilcox test to compare the means of fermentation alpha-diversities.

### Monitoring of inoculated starters in fermentation

To confirm the starters added in the defined times were present during fermentations (F1, F2 and F3) the *16S rRNA* and *ITS* genes sequenced by the Sanger method were used as reference in a blast analysis to correlate the ASVs *16S rRNA* reads with their cognate sequence. Yeasts DNA was extracted from two-days incubated cultures grew on YPD medium. From cultures, an aliquot of 2.0 ml was selected, and the DNA was obtained using the Wizard Genomic DNA Purification Kit (Promega, Milan, Italy), following the manufacturer’s instructions. DNA concentration was determined using a Nanodrop®. *ITS* 5.8S *rRNA* gene was amplified using the primers ITS1 (TCCGTAGGTGAACCTGCGG) and ITS4 (TCCTCCGCTTATTGATATGC) (White, 1990). The PCR reaction was performed as described by Esteve-Zarzoso et al. (1999). Finally, PCR products were purified using the kit Gene elute^TM^ Gel extraction kit (Sigma Aldrich), re-surveyed on Nanodrop® and delivered for Sanger DNA sequenced at Eurofins Genomics (Ebersberg, Germany).

LAB DNA was extracted from pure cultures grew overnight. From the culture, an aliquot of 1.0 ml was selected and centrifuged at 14,000 rpm at 4°C for 5 min. The underneath was discarded, and the pellet was pre-treated for disruption of the bacterial cell wall with 300 μl of lysozyme solution in Tris-EDTA buffer (TE) (10 mg/ml, Sigma-Aldrich/Merck), incubated at 37°C for 60 min. The resulting suspension was then centrifuged at 14,000 rpm at 4°C to remove cellular fragments and the genomic DNA finally was extracted using the Wizard® Genomic DNA Purification Kit reference A1125 (Promega, Madison, WI, USA) following the bulla guidelines. The pure DNA was quality-assessed and quantified using Nanodrop® (Thermo Fisher Scientific, USA). Full *16S rRNA* gene was amplified using the primers E8F (5’AGAGTTTGATCCTGGCTCAG3’) and E1541R (5’AAGGAGGTGATCCANCCRCA3’) (Baker et al., 2003). PCR was performed as described by Rathnayake et al. (2010). The PCR products were purified using the kit Gene elute^TM^ Gel extraction kit (Sigma Aldrich, MA, USA), re-surveyed on Nanodrop® and delivered for Sanger DNA sequenced at Eurofins Genomics (Ebersberg, Germany).

*A. senegalensis* strains’ DNA was extracted from overnight cultures on BHI medium (OXOID, Basingstoke, UK) from a 1,0 ml aliquot of suspension was selected for DNA extraction. The DNA was extracted using the kit Illustrative Bacteria Genomic Mini Spin Kit (GE Life Sciences, Switzerland) according to manufacturer’s recommendations. The pure DNA was quality-assessed and quantified using Nanodrop® (Thermo Fisher Scientific, USA). Full *16S rRNA* was amplified using the primers 27F (5’AGAGTTTGATCMTGGCTCAG3’) and (5’GCTTACCTTGTTACGACTT3’) according to the protocol of Tulini et al. (2016). The amplicons were visualised on an agarose gel (0.8%) electrophoresis, and amplicons were purified using the Gel Band Purification kit (GE Life Sciences), re-surveyed on Nanodrop® and delivered to DNA Sanger sequencing at “laboratório de sequenciamento de ácidos nucleicos da FCFRP-USP” (Brazil). Bacterial and fungal sequencing amplicons were made publicly available on the GenBank database. Accession numbers ON623883-ON623889 and ON738686-ON738691, respectively.

A threshold of 97% of identity was established as minimal to assign an ASV to the respective species based on *16S rRNA* and *ITS* gene. The sequences presenting alignments with no more than 97% of similarity were blasted against the NCBI nr-database in order to relate the ASV with a cognate species.

### Statistical analysis of correlation and regression

Correlation among ASVs and measured variables (i.e., volatile compounds and *luxS* gene expression) was performed using the R package microbiomeSeq (Ssekagiri et al., 2017). Regression analyses among *luxS* gene expression versus LAB loads was performed using the R package ggpmisc (Aphalo, 2020).

## Results

In this study, four lab-scale fermentations were performed to unravel whether the influence of quorum sensing on microbial shifts has possible impacts on sensorial attributes development along cocoa fermentation. The experiments were planned to aim to understand the interaction of the autochthonous microbiota and selected starter cultures employed, as the raw material was not sterile to simulate field conditions. All fermentations exhibited microbial successions as determined by microbial enumeration and NGS. In terms of pH and temperature, no significant differences were observed for these variables, being the pH variation of 3.30 up to 3.45 for the experimental fermentations and 3.2 up to 4.5 for the non-inoculated one (F0). Moreover, temperature was not different among fermentations, reaching a maximum of 26°C in laboratory conditions (Supplementary Fig. A.1). Regarding microbiota, microbial loads presented by experimental fermentations (F1, F2 and F3) were similar to the F0 fermentation as shown in Fig. 2A and were not differentially significant (Wilcoxon test p-value > 0.05). Besides, NGS corroborated enumeration analysis, revealing no significant differences (p-value > 0.05) in α-diversity dynamics caused by starters inoculation (Fig. 2B). Nonetheless, F1 and F3 fermentations presented augmented median values of observed and expected (Chao1) indexes, meaning these fermentations were more diverse in terms of intrasample microbial composition (a slightly enhanced number of different microorganisms detected). In contrast, in terms of equitability (Shannon measurements), F2 fermentation presented lower variations (boxplot heights). In addition to the observed number of ASVs and Chao1 estimates, the equitability of F2 fermentation suggests a low variation in microbial diversity and composition over the fermentation period (Fig. 2B). In summary, even with the addition of defined cocktails in F1, F2 and F3 fermentations, in general it was observed a restricted microbial diversity, which not necessarily means the composition was not impacted by the addition of starter cultures, but fermentations were not dissimilar regarding changes in microbial richness.

**Fig. 2.**
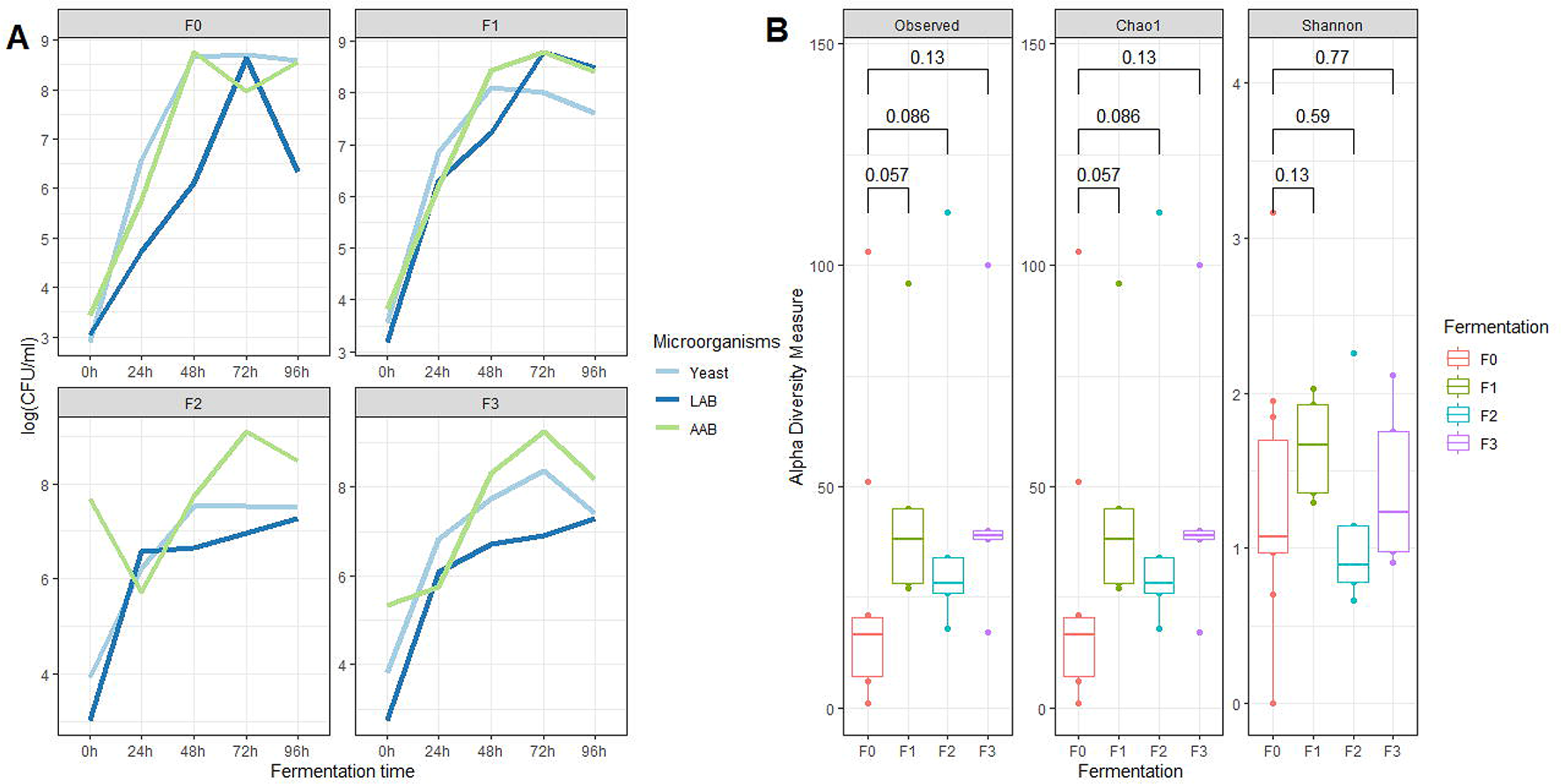
(A) Enumeration of culturable microbial groups in fermentations and (B) representation of uncultured microbial diversity generated by NGS. The comparison lines above box plots and respective numbers stands for p-values calculations of Wilcoxon test from the comparison of F0 and F1, F2, and F3 fermentations.

### Microbial composition was partially influenced by starters

While discrete fluctuations of microbial diversity and low diversity exhibited by experimental fermentations, the inoculation of different cocktails resulted in distinct bacterial profiles, except for F2 and F3 which presented a slight semblance. In comparison to the non-inoculated fermentation (F0), characterised by a heterogeneous composition of non-dominant taxa (glued as <0.01% of representativity) (Fig. 3A), F1 fermentation exhibited a dominance of LAB, specifically lactobacilli (older genus *“Lactobacillus”*, Zheng et al. 2020) in the middle to the last phases of fermentation (48h-96h) (Fig. 3A). As in this fermentation all cocktails (composed by yeasts, LAB, and AAB) were inoculated, it was expected to observe higher amounts of AAB bacteria in the range of 72h-96h of fermentation, which was observed only in the final period of fermentation. F2 and F3 fermentations were characterised by the dominance of *Pantoea* (enterobacteria) from the middle to the last phases (48h-96h) and AAB at 96h of fermentation (Fig. 3A). The presence of AAB in the F2 fermentation was expected since only with the C3 cocktail was inoculated in that fermentation, but for the F3 fermentation the AAB dominance was unexpected as only the C1 cocktail was added. Another interesting observation was the ability of all cocktails (C1, C2, and C3) to replace accompanying detrimental microbiota, such as enterobacteria which was detected only in the start of fermentations (0h) for the inoculated fermentations (F1, F2, and F3) (Fig. 3A). Besides, *Gluconobacter* was detected after 72h of fermentation in all experiments. ASVs related to this genus were also detected in F0 fermentation (Fig. 3A).

**Fig. 3.**
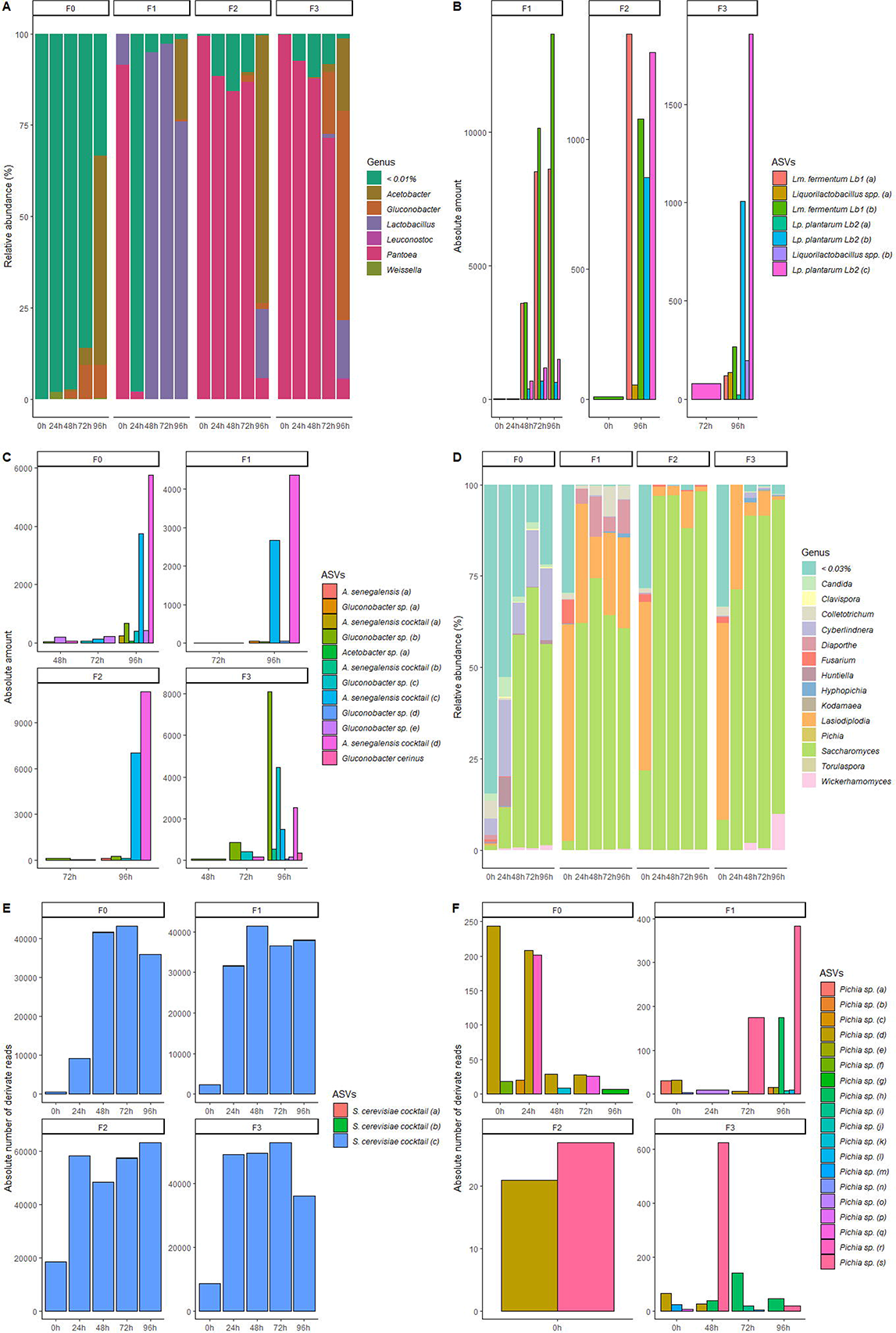
Microbial composition along cocoa fermentation. (A) Bacterial composition of main genera detected along fermentation. (B) Scrutinization of ASVs identified as lactobacilli and association with inoculated strains (determined by blasting ASVs against strains’ *16S rRNA* gene) and autochthonous ones (determined by blasting ASVs against the NCBI database). (C) Depiction of ASVs related to *Acetobacter* sp. and association with inoculated strains (determined by blasting ASVs against strains’ *16S rRNA* gene) and autochthonous ones (determined by blasting ASVs against the NCBI database). (D) Fungal composition of main genera detected along fermentation. (E) Representation of *S. cerevisiae* ASVs diversity in fermentation and association with inoculated strains (determined by blasting ASVs against strains’ *16S rRNA* gene) and autochthonous ones (determined by blasting ASVs against the NCBI database). (F) Depiction of ASVs related to *Pichia* genus and association with inoculated strains (determined by blasting ASVs against strains’ *16S rRNA* gene) and autochthonous ones (determined by blasting ASVs against the NCBI database).

Regarding the monitoring of starters fluctuations along fermentations, the taxonomic assignment for this level was satisfactory for the identification of two ASVs derived from the *Lm. fermentum* Lb1 strain, which dominated the middle to the last phases of F1 and were detected also in F2 (96h) and F3 (96h-lower amounts) fermentations (Fig. 3B). The *Lp. plantarum* Lb2 strain was related to three ASVs, being the ASVs “b” and “c” enriched in the F2 and F3 fermentations, which were not inoculated with the C2 cocktail. Other lactobacilli-related ASVs were classified by blast analysis against the NCBI nr-database since they do not present similarity with *16S rRNA* gene of the strains inoculated in fermentations. These lactobacilli were classified only in the genus level as *Liquorilactobacillus* spp (Fig. 3B). Moreover, it was not detected as lactobacilli ASVs in the control (F0) fermentation.

As expected, due to the latter inoculation (48h) of the C3 cocktail, *Acetobacter* spp. were detected in the final of fermentations (96h) along with *Gluconobacter* spp (Fig. 3A). The addition of a defined cocktail of *A. senegalensis* strains (C3-cocktail) was not enough to allow *Acetobacter* dominance in F1 and F2 fermentations, as this genus was also determined in the last phases of F0 and F3 fermentations as well (Fig. 3A). The monitoring of the inoculated strains through the similarity of *16S rRNA* genes and the ASVs sequences was achieved and demonstrated the presence of six *Acetobacter* spp. ASVs, being four of them, originated from the inoculated *A. senegalensis* strains (Fig. 3C). However, no differentiation of ASVs originating from a cognate strain was possible as blast analysis evidenced a high similarity among the *16S rRNA* genes of these strains. Besides, some ASVs identified as *Acetobacter* sp. on Qiime2 were identified as *Gluconobacter* sp. by blast analysis against the NCBI nr-database (Fig. 3C). Only the ASV “c” belonging to the inoculated *A. senegalensis* strains was detected in significant amounts at the period of 96h of fermentation in most experimental fermentations (Fig. 3C).

The taxonomic composition of fungi was more diverse than bacteria and varied among fermentations. F0 fermentation was dominated by ASVs related to the *Saccharomyces* genus from the middle to the end of fermentation (48h-96h), while the inoculated fermentations presented a robust dominance of this genus along the entire time range (Fig. 3D). F1, F2, and F3 fermentations presented higher amounts of *Lasiodiplodia* at the start, being the middle and the end of fermentations marked by *Saccharomyces* dominance. *Diaporthe* ASVs were observed in the range of 24h-96h in F1 fermentation in association of *Candida* ASVs fluctuations in the period 0h-96h (Fig. 3D). The remaining F2 and F3 fermentation were virtually equivalent in terms of fungal composition.

Concerning the fungal species of interest (whose genera *Saccharomyces* and *Pichia* were inoculated in fermentations), only three ASVs were assigned to *Saccharomyces* genus and to the *S. cerevisiae* species, being impossible to distinguish from which strains these sequences were from (Fig. 3E). Only the “c” ASV was dominant in all fermentations, while the ASVs “a” and “b” were detected in low amounts (Fig. 3E). Concerning *Pichia*, taxonomic assignment was unable to assign the *Pichia* ASVs to the species level (Fig. 3F). Besides, *Pichia* ASVs exhibited a huge diversity of ASVs, summing at least 19 amplicon variants with uneven prevalence in fermentations (Fig. 3F).

### *Lp. plantarum* species trend to express more *luxS* than *Lm. fermentum*

The Fig. 4 presents the data related to the expression of the *luxS* gene throughout the evaluated fermentations. Regarding the *luxS* gene expressed by the species *Lp. plantarum* (Fig. 4A), it was observed that the addition of the C2 cocktail (composed only by the strains *Lp. plantarum* Lb2 and *Lm. fermentum* Lb1) was able to enrich the expression of this gene in the F1 fermentation from the middle (48h) to the end (96h) of the fermentation. In relation to 48h of fermentation, the increase in gene expression was about 66.67 times greater than in the same period determined for F0 fermentation. Gene expression was linearly increasing at 72h and 96h fermentation times, which in relation to F0 fermentation was about 1,250 and 166.67 times greater, respectively. For the F2 and F3 fermentations, the expression of the *luxS* gene species-specific of *Lp. plantarum* was detected in the range of 72h and 96h of fermentation, being higher than the F0 fermentation only at the end of fermentation (96h) (Fig. 4A). During this period, the F2 and F3 fermentations showed higher expression of *luxS* related to the *Lp. plantarum* species are about 2.67 and 6.67 times greater, respectively, in relation to the F0 fermentation in the same period (Fig. 4A).

**Fig. 4.**
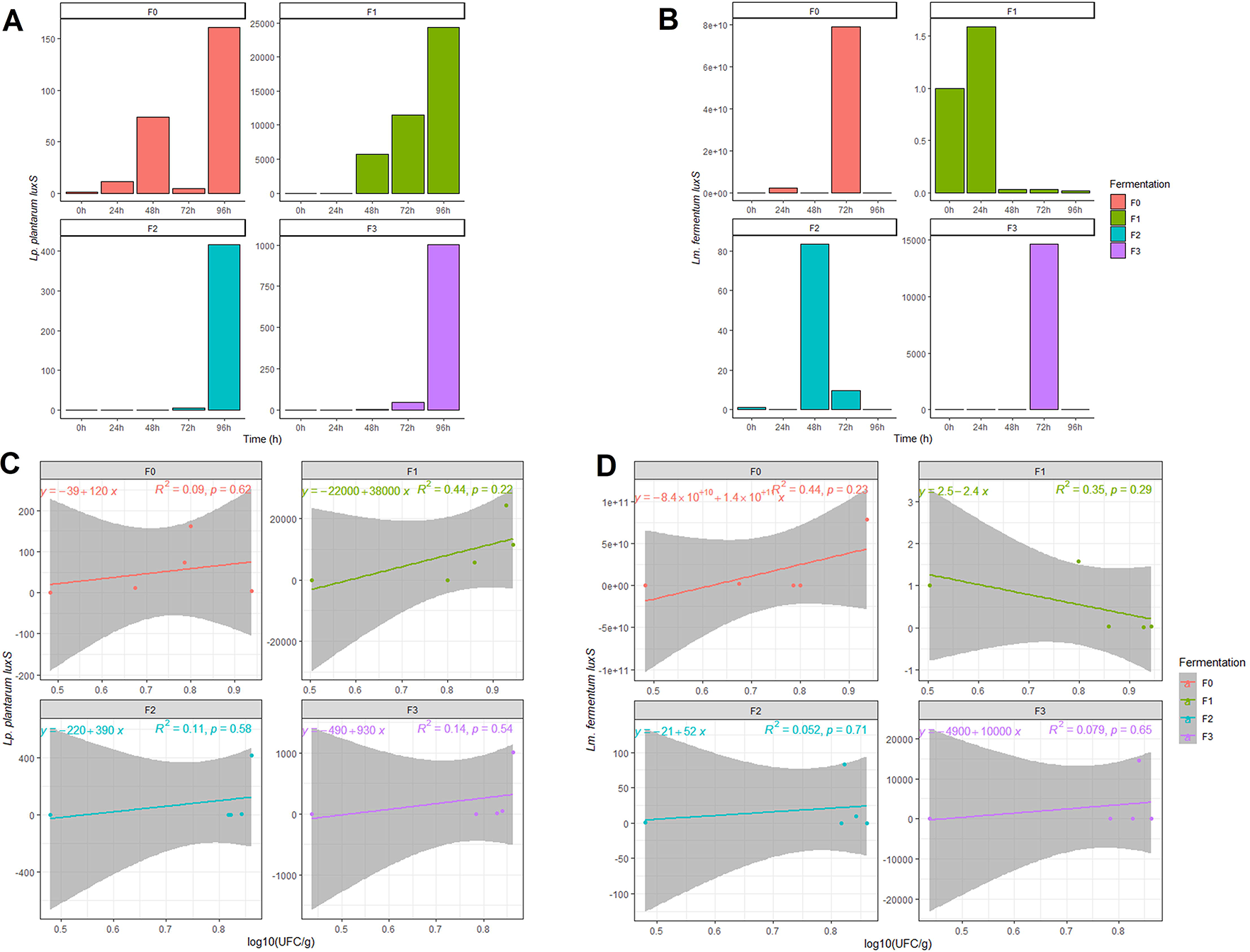
*luxS* gene expression in lab-scale cocoa fermentations. (A) Variation of 2^-ΔΔCt^ values of *Lp. plantarum luxS* gene expression along fermentation. (B) Variation of 2^-ΔΔCt^ values of *Lm. fermentum luxS* gene expression along fermentation; (C) Linear regression of *Lp. plantarum luxS* gene expression versus LAB enumeration (cellular density) along fermentation; (D) Linear regression of *Lm. fermentum luxS* gene expression versus LAB enumeration (cellular density) along fermentation.

Curiously, the expression of the *luxS* gene by the species *Lm. fermentum*, even with the addition of the C2 cocktail, was not higher in relation to the F0 fermentation (Fig. 4B). Furthermore, only in F0 fermentation the expression of this gene was enormously increased but detected in large amounts only in the 72h period of fermentation (Fig. 4B). According to the data presented, *Lp. plantarum* tends to show greater expression of the *luxS* gene than the species *Lm. fermentum*. However, no pattern could be observed since there was no significant difference (Wilcoxon Test p-value>0.05) among fermentations in terms of gene expression. Furthermore, even comparing the density of lactic acid bacteria over time (presumptive LAB plate enumeration data) with the increase in *luxS* gene expression (Figs. 4C and 4D), in all fermentations as LAB cell density increases, there is an increase in *luxS* gene expression of both lactobacilli species, except for F1 fermentation, in which *luxS* expression by *Lm. fermentum* was inversely related to the increase in cell density.

### Metabolic changes along fermentation and production of volatile compounds

Enzymatic degradation of cocoa pulp is crucial for flavour development as the infiltration of external compounds in consonance with endogenous hydrolases lead to a multitude of VOCs which directly impact sensorial attributes of fermented seeds (Aprotosoaie et al., 2016; Batista et al., 2016). In this study, 16 enzyme families were dosed: amilases, β-glucanases, cellulases, arabinofuranosidases, invertases, endoglucanases, xylanases, β-glucosidases, arabinases, xyloglucanases, esterases, β-xylosidases, pectinases, mannanases, cellobiohydrolases, and lipases (Fig. 5).

**Fig. 5.**
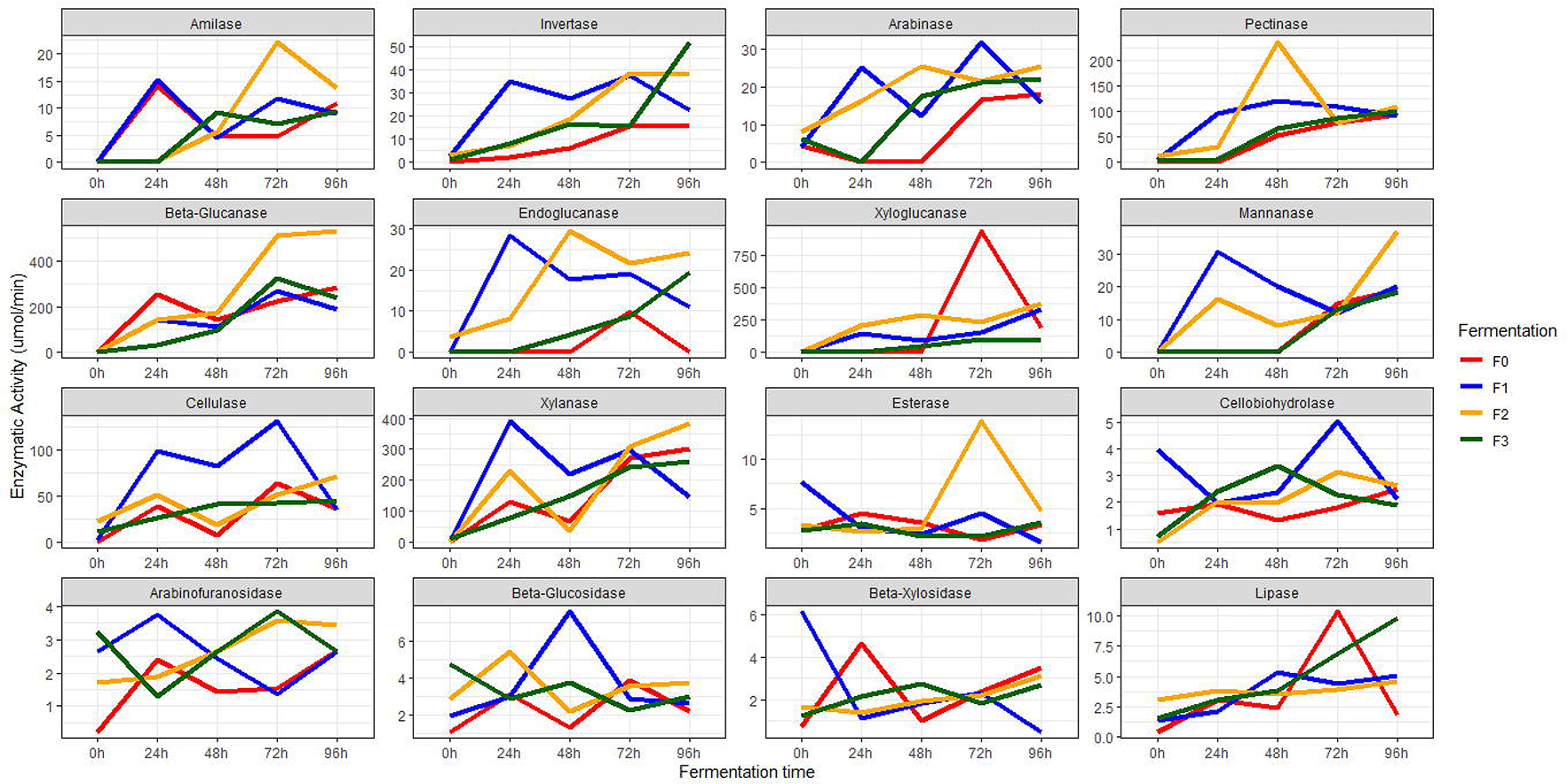
Enzymatic profiles measured in surveyed fermentations related to complex carbohydrates and polysaccharides breakdown.

Higher enzymatic activities were determined for cellulases (72h), arabinofuranosidases (24h), invertases (24h), endoglucanases (24h), xylanases (24h), β-glucosidases (48h), arabinases (72h), mannanases (24h), and cellobiohydrolases (72h) in F1 when compared to the F0 fermentation. Amilases (72h), β-glucanases (72h), arabinofuranosidases (72h), invertases (72h), endoglucanases (72h), arabinases (48h), esterases (72h), and pectinases (48h) were augmented in F2 in relation to F0 fermentation. Arabinofuranosidases (72h), invertases (96h), arabinases (72h), and cellobiohydrolases (48h) were higher in F3 fermentation than F0. Interestingly, lipase activity was pronounced in F0 and F3 fermentations, and xyloglucanase activity was higher in the control fermentation than others (Fig. 5).

Additionally, the higher enzymatic activity determined in F1, F2 and F3 fermentations were proportional to the microbial shifts caused by inoculation of starter cultures. In F1, for instance, the augmented activities of cellulases (72h), arabinofuranosidases (24h), invertases (24h), endoglucanases (24h), xylanases (24h), β-glucosidases (48h), arabinases (72h), mannanases (24h), and cellobiohydrolases (72h) coincided to the peak of LAB dominance in that fermentation period (24h-72h), while in F2 the prevalence of higher enzymatic activities in the range of 48h-72h coincided to the period of AAB inoculation (Fig. 5).

To track the possible microbial actors related to enzymatic changes in cocoa seeds along fermentation a Kendall correlation was performed among genera and measured enzymatic activities. Few genera of the inoculated strains were significatively correlated to enzymatic changes (Fig. 6). *Pichia* and Lactobacilli were negatively associated with invertases and pectinases in F0, while for the experimental fermentations no significant association was observed (Fig. 6). Nevertheless, even with no statistical meaning, it’s worth to emphasise that in most fermentations *Pichia* was positively correlated with esterase activity (F0 and F1), while *Saccharomyces* was positively correlated with amilase (F1 and F2), arabinase (F1), β-glucosidase (F1), cellulase (F1), cellobiohydrolase (F2 and F3), endoglucanase (F1), invertase (F1), mannanase (F1), pectinase (F1) and xylanase (F1). Lactobacilli exhibited higher correlation to arabinase (F3), β-glucanase (F1), endoglucanase (F1) and mannanase (F3). *Acetobacter* instead was more related to arabinose (F0), invertase (F0), mannanase (F0 and F3), pectinase (F0), β-xylosidase (F2) and esterase (F2) (Fig. 6).

**Fig. 6.**
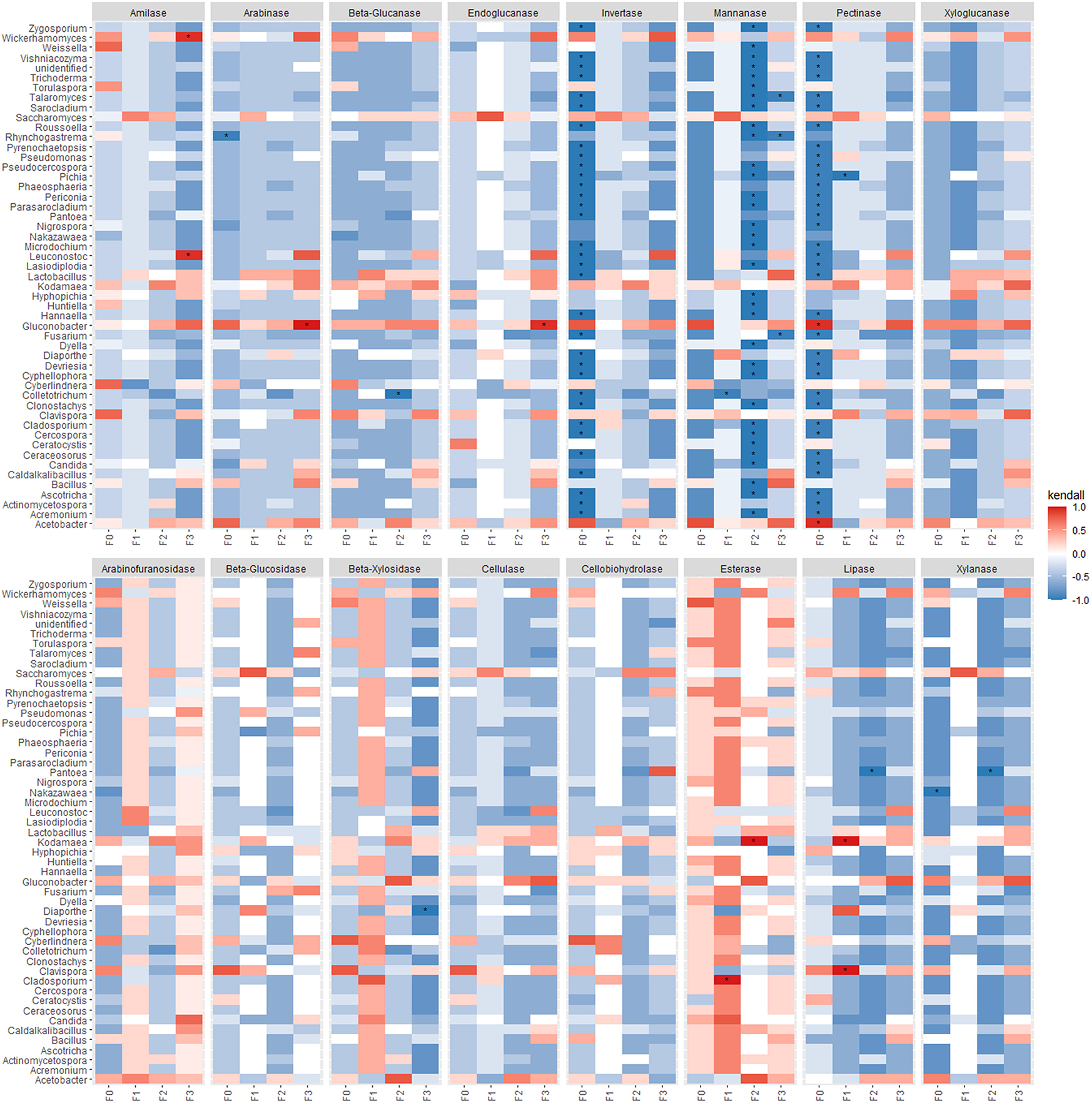
Correlation of enzymatic profiles and microbial composition determined by the Kendall correlation test.

The results of microbial metabolic activities were not only measured by enzymatic ability to breakdown complex sugars, but by the production of VOCs as well. Table 2 describes the VOCs identified by a qualitative HPLC analysis in both replicates of each inoculated fermentation (F1, F2 and F3) and the control one (F0). A total of 89 compounds were identified. Most of them representing alcohols and phenols (n=28), aldehydes and ketones (n=25), esters (n=13), and sulfur compounds (n=3) (Table 2) distributed unequally in fermentations. 20 compounds were not identified as playing a significative role, based on literature reviewing (Aprotosoaie et al., 2016; Misnawi & Ariza, 2011; Owusu et al., 2010; Rodriguez-Campos et al., 2011), on flavour production in cocoa fermentation, and so were represented by a trace in the Table 2.

**Table 2.**
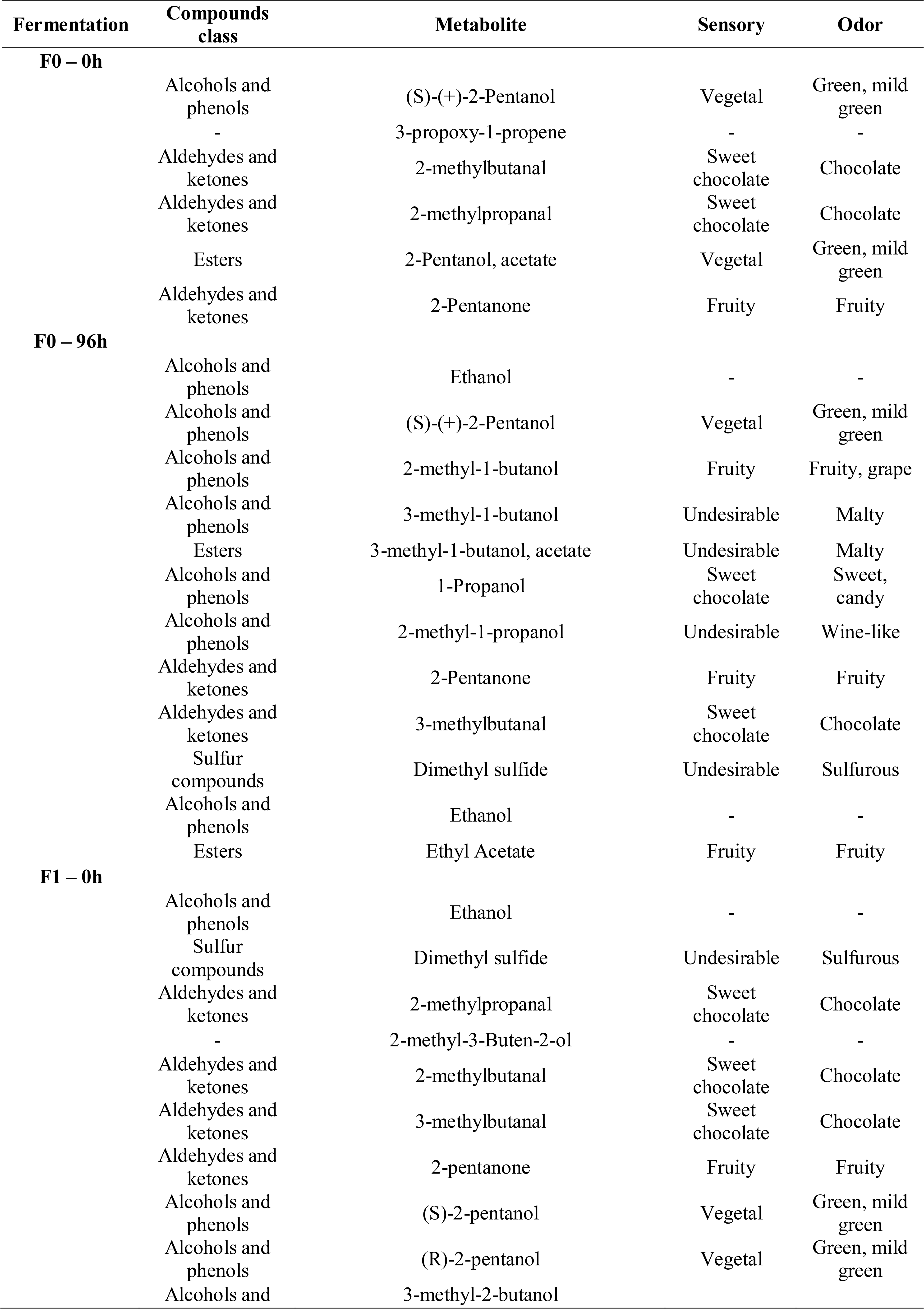
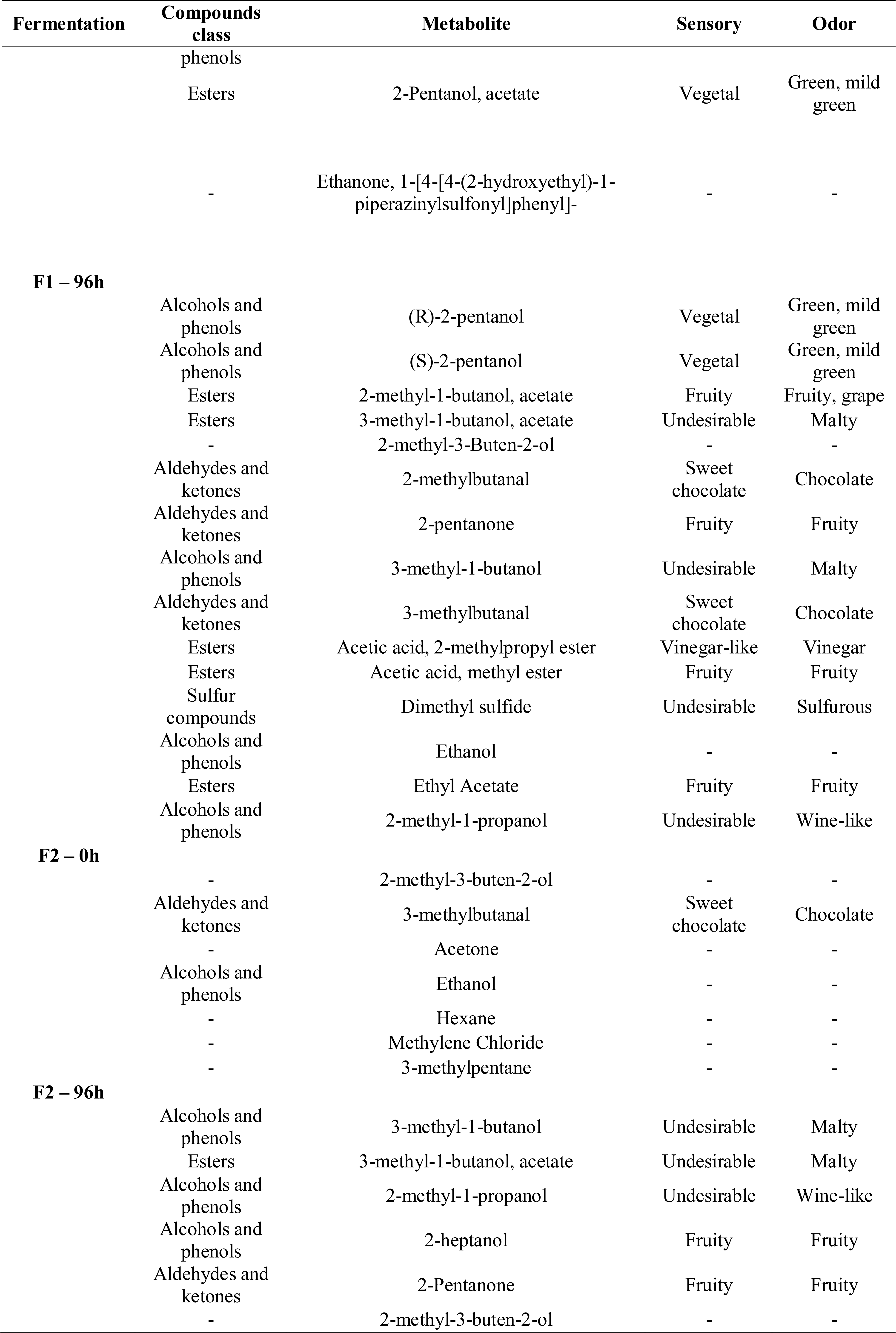
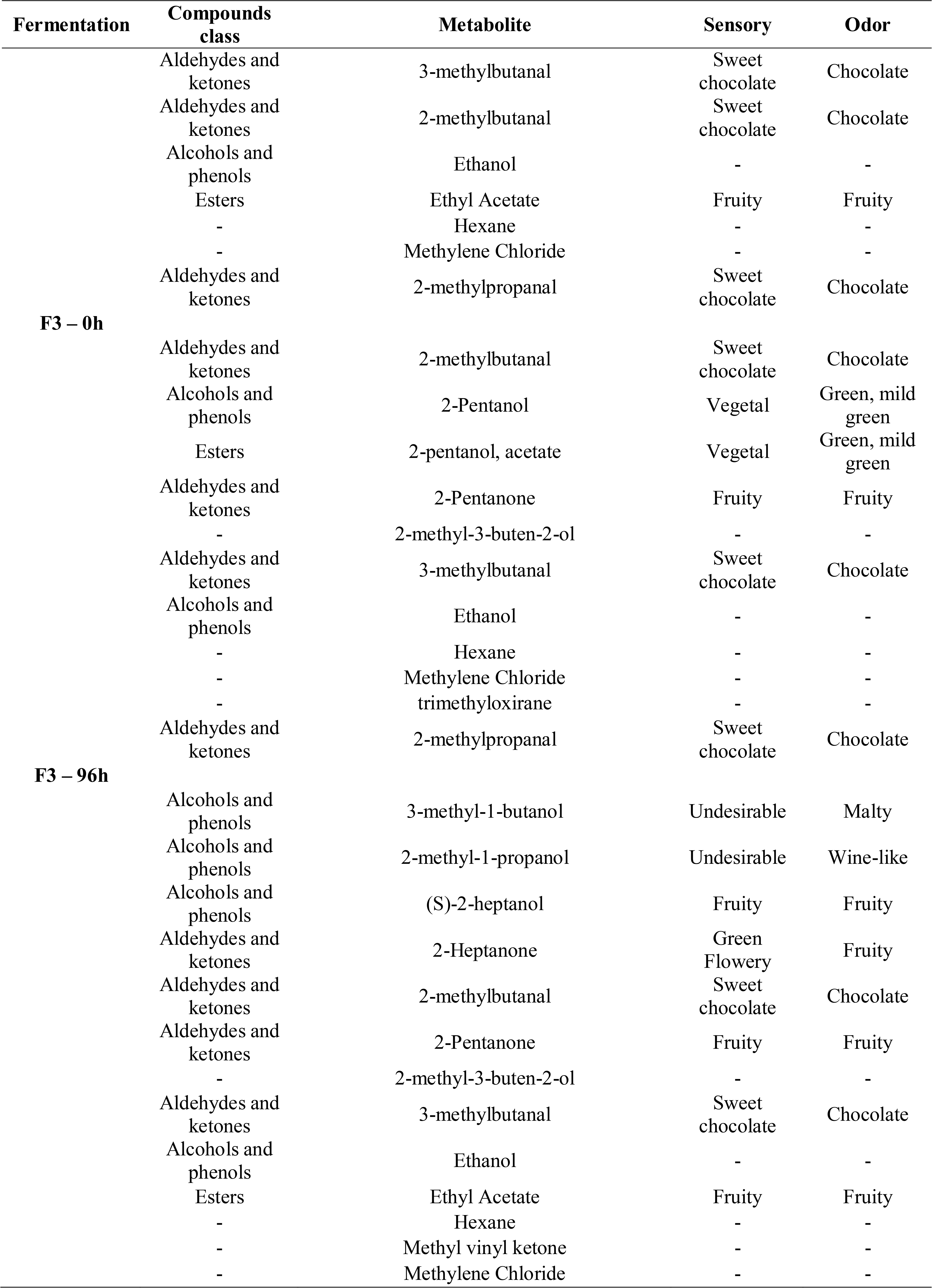
Detailing of volatile metabolic compounds detected in experimental fermentations and sensorial characteristics.

Fig. 7 shows the scores of surveyed fermentations and demonstrates that even with the addition of defined cocktails the risk of generation of detrimental flavour precursors occurs. In all fermentations, but F2, is possible to visualise a tendency for sweetness-, fruity- and vegetal-like VOCs precursors at the beginning of fermentations, while at the end of the processes the emergence of undesirable compounds, such as wine-like and malty odors were detected. Nevertheless, some desired sensorial attributes were exalted. For example, in F0 fermentation the detection of undesirable compounds was balanced by the augment of fruity and chocolate sweetness precursors (Fig. 7). In F1 instead, at the end of fermentation, there was a reduction of chocolate notes, emergence of undesirable and vinegar-like VOCs, with the concomitant increasing of fruity notes. In F2 the tendency of production of undesirable VOCs was counterbalanced by the enhancement of fruity- and chocolate-like VOCs (Fig. 7). In F3 fermentation, there was an augment of undesirable compounds, decreasing of chocolate-like precursors and an enhancement of fruity-like VOCs (Fig. 7). Statistical analysis based on the Wilcoxon test showed the differences observed in the start to the end point of fermentations were not significant (p-value > 0.05).

**Fig. 7.**
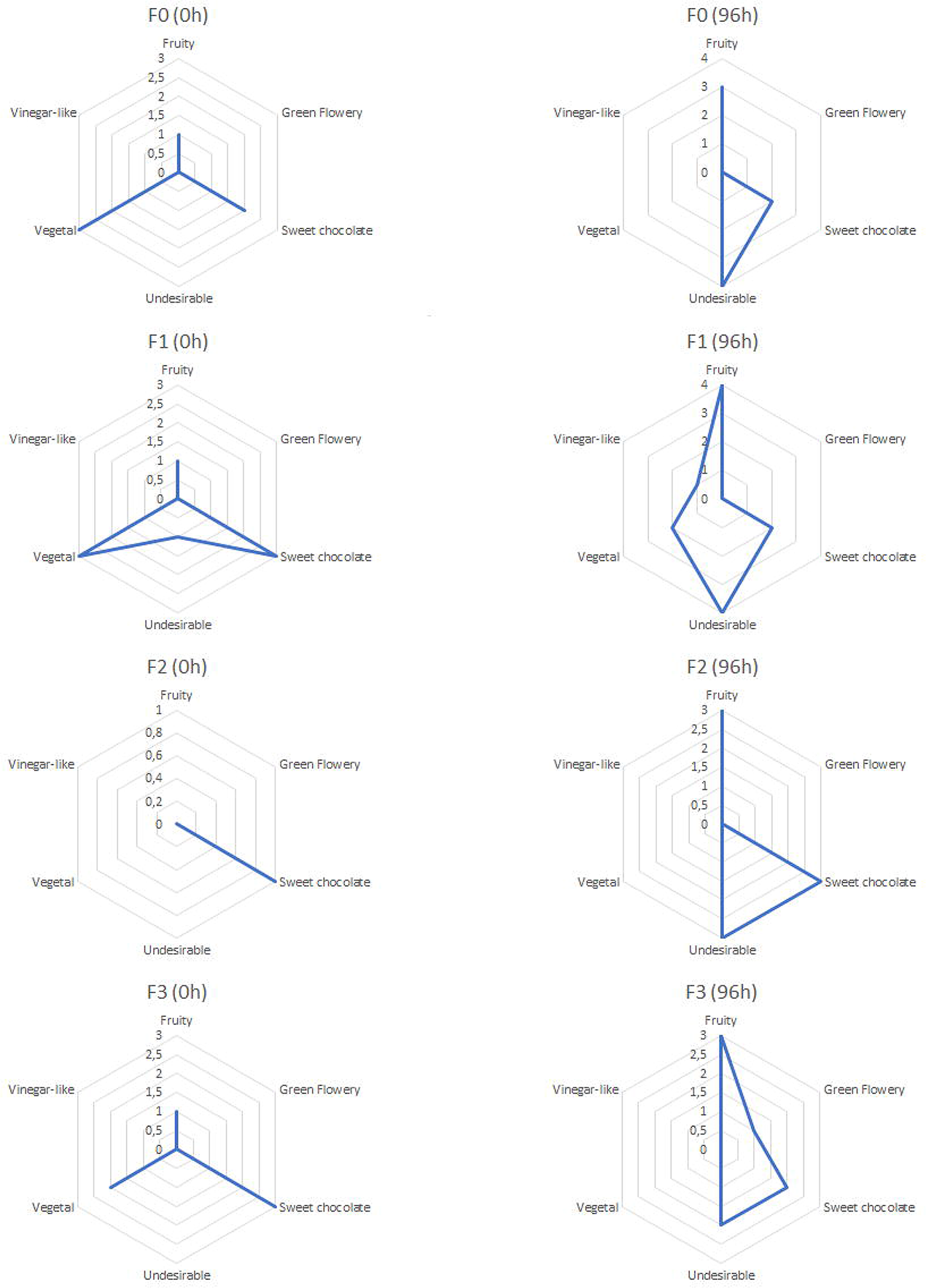
Radar plots representing the scores of sensorial traits present at the beginning (0h) and at the end (96h) of fermentations.

In order to infer whether QS has a potential contribution for cocoa fermentation quality a Kendall correlation was performed among *luxS* expression amounts (at the beginning and end of fermentations), and the qualitative presence/absence of sensorial precursors determined by HPLC (at the beginning and end of fermentations) (Supplementary Fig. A.2). From the data derived from this study, it was impossible to precisely determine if QS directly impacted the quality of fermentations as no significant correlation (p < 0.05) was observed among *luxS* gene expression and sensorial profiles. However, the production of some precursors such as vegetal-, vinegar-, fruity-like (this presenting p < 0.05) and undesirable (i.e., malty- and wine-like) compounds was correlated with *Lp. plantarum luxS* gene expression. Conversely, Vegetal-like notes were more associated with *Lm. fermentum luxS* expression (Supplementary Fig. A.2).

## Discussion

Cocoa fermentation has a huge importance for the development of sensorial traits in the raw material for chocolate’s production. Several studies have been conducted to propose novel starter cultures to standardise the process, relating the dominance of candidate strains with the development of sensorial attributes along fermentation (Batista et al., 2016; García-Ríos et al., 2021; Magalhães da Veiga Moreira et al., 2017; Mota Gutierrez et al., 2019; H. G. Ouattara et al., 2020). Following this direction, the present research aimed to investigate the possible contribution of QS for bacterial dominance, specifically lactic acid bacteria, in fermentation and if this could result in fermented beans with superior quality in terms of production of aroma precursors.

To scrutinise this hypothesis, yeasts, LAB, and AAB isolates from a spontaneous cocoa fermentation (Almeida et al., 2020) were selected as they are intrinsic to the cocoa and fermentation ecosystem. The yeasts *S. cerevisiae* strains F3, F6, F11 and F12 and the strains *Pichia kudriavzevii* PCA1 and *Pichia kluyveri* strain PCA4 were identified and genotypically characterised in this work, while the LAB *Lm. fermentum* Lb1 and *Lp. plantarum* Lb2, the AAB *A. senegalensis* strains MRS7, GYC10, GYC12, GYC19 and GYC27 were previously evaluated and genomic characterised (de Almeida et al., 2021; Almeida et al., 2022). These yeasts were selected to add contrast of fungal diversity and due to the valuable importance of fungi for cocoa fermentation (Crafack et al., 2013; Lefeber et al., 2012; Magalhães da Veiga Moreira et al., 2017), while the LAB and AAB were selected given the QS potential teased previously (Almeida et al., 2021) and adaptive potential to stressors (Almeida et al., 2022), respectively.

Cocoa fermentation is hard to standardise, but several studies emphasise the dominance of *S. cerevisiae*, *Lp. plantarum*, *Lm. fermentum* and *Acetobacter* species as core members (Lefeber et al., 2011; Meersman et al., 2013; Papalexandratou et al., 2011; Verce et al., 2021), with evidence to *Lm. fermentum* to the detriment of *Lp. plantarum*. In this work we employed a deep marker analysis based on the monitoring of ASVs along fermentation to assess whether the inoculated strains participated or not in fermentations. However, the technology of ASVs determination is not always able to distinguish among closely related strains but refers to the truly biological meaning of a sequence variant. In other words, ASVs can separate sequencing errors from biological variations in sequences (Callahan et al., 2017). In cocoa fermentation, ASVs were already employed by oligotyping methods (ASVs), which revealed a huge intraspecific variability of yeasts and bacteria (LAB and AAB) along fermentation (Díaz-Muñoz et al., 2021; Pacheco-Montealegre et al., 2020; Verce et al., 2021). Therefore, to discriminate if the ASVs might represent intraspecific strains variability or if a given strain originated several ASVs, the comparison of gene markers (*16S rRNA* and *ITS*) against the ASVs provides a useful resolution (Verce et al., 2021; Díaz-Muñoz et al., 2021).

The inoculation of *Pichia kudriavzevii* strain PCA1 and *Pichia kluyveri* strain PCA4 produced no significant changes in fermentations (F1, F2 and F3), as these strains were not dominant and no *Pichia* ASVs found matched the inoculated ones, suggesting these strains cannot replace indigenous *Pichia* species. The high intraspecies diversity of *Pichia* spp. (H. G. Ouattara & Niamké, 2021; Pereira et al., 2017) could explain the high numbers of different ASVs measured in this study. Regarding *S. cerevisiae*, it was shown that the inoculated strains were dominant in F1, F2 and F3 fermentations, but it was impossible to distinguish which strain dominated the scenarios as they presented high similarity in their *ITS* gene sequences. As observed, even without inoculation of any cocktail, the F0 fermentation presented the same *S. cerevisiae* strains, suggesting these strains are indigenous strains present in cocoa producing regions of Brazil and could be used for starter development. These observations are in accordance with the literature, as *Saccharomyces cerevisiae* strains trend to dominate fermentations from different regions and *Pichia* sp., especially *Pichia kudriavzevii* are more variable in genetic background, being dispersed in several regions (Ouattara et al., 2021).

Although no statistical difference was observed among the experimental fermentations, the F1 fermentation presented the highest loads of ASVs derived from *Lm. fermentum* reads in comparison to the *Lp. plantarum* ones. The *Lm. fermentum* dominance in cocoa fermentation is not novel and it was already explained in terms of physiological adaptations. It was demonstrated that heterofermentative *Lm. fermentum* isolates obtained from cocoa fermentation (indigenous strains) may be able to metabolise citric acid, oppositely to the type strain, as this characteristic seems species-specific and corresponds to the reality in the cocoa ecosystem: the pulp is rich in citric acid (Lefeber et al., 2010).

In cocoa fermentation, *Lp. plantarum* and *Lm. fermentum* species preferentially consume fructose in relation to glucose. This leads to mannitol reduction and acetic acid production by a heterofermentative metabolism. Thus, the metabolization of citric acid aids the dominance of *Lm. fermentum*, which could augment the quality of fermentation by replacing non-citrate fermenters (depectinizing yeasts), which capture voraciously carbohydrates and convert them directly to ethanol. Therefore, more production of acetic acid from fructose consumption and mannitol reduction allows the regeneration of NAD+ for extra ATP production, aiming for bacterial proliferation (Lefeber et al., 2010). Nevertheless, lactobacilli metabolism is inhibited in acid titles above 5%, limiting their growth and colonisation (Ouattara et al., 2016).

Regarding the temporal distribution of *Lp. plantarum* (beginning-middle of fermentation) and *Lm. fermentum* (middle-end of fermentation), surveys of spontaneous fermentations have already observed the shifts displayed by these species (Camu et al., 2007; Papalexandratou et al., 2013). As observed for *S. cerevisiae* strains, the same *Lp. plantarum* Lb2 and *Lm. fermentum* Lb1 strains were detected in the F0 fermentation, suggesting these strains are dominant in cocoa producing regions.

However, some lactobacilli ASVs were identified as *Liquorilactobacillus* spp, a genus commonly found in cocoa fermentation (Almeida & De Martinis, 2021). As the novel taxonomic classification was proposed (Zheng et al., 2020), the older genus *“Lactobacillus”* was now split in several genera, among them *Limosilactobacillus*, *Lactiplantibacillus*, and *Liquorilactobacillus*. The huge intra-genera variability explored by Zheng et al. (2020) explains the reason of previously classified *“Lactobacillus”* species on SILVA database were identified as *Liquorilactobacillus* spp. in this work, as to confirm the species level assignment, each ASV was blasted against the *16S rRNA* genes of the starter cultures strains and against the NCBI non-redundant database, which is already updated for the novel taxonomy. So, as reported by a metagenome-assembled genomes survey (Almeida & De Martinis, 2021), many microorganisms may still be overlooked in cocoa fermentation, and the new taxonomy sheds light into the huge variability of previous taxa which were named “*Lactobacillus”*, but in fact belong to other LAB genera.

*Acetobacter* species are dominant in cocoa fermentation (Miescher Schwenninger et al., 2016) and *A. senegalensis* was already proposed as a potential starter candidate (Illeghems et al., 2016). In this work, four of six *A. senegalensis* ASVs identified matched to the inoculated strains but no differentiation between the strains from C2 cocktail was possible due to higher similarities in their *16S rRNA* gene sequences. The determination of *A. senegalensis* strains was masked by the ambiguous identification of *Gluconobacter* species presenting high similarities for the *16S rRNA* gene as, during blast analysis, some *Acetobacter* ASVs were identified as *Gluconobacter* spp. However, the results raised in this study show no dominance of the inoculated *Acetobacter senegalensis* strains, which were detected in lower amounts, even presenting an interesting metabolic potential to circumvent adverse conditions in fermentative environments (Almeida et al., 2022).

### Metabolization of cocoa seeds and VOCs production

To the best of the authors’ knowledge, no study was still performed to measure enzymatic activities directly in cocoa fermentation and correlate it to microbial dynamics. In the present research, no significant association was made regarding microorganisms and enzymatic activities, which suggests the main enzymatic profiles determined derivate from the pulp and cotyledons metabolism (endogenous hydrolases) as disseminated in literature (De Vuyst & Weckx, 2016). A study showed the potential of some yeasts strains isolated from cocoa fermentation to produce β-glucosidases, xylanases, pectinases, cellulases, and lipases (Delgado-Ospina et al., 2020). Another study demonstrated the influence of *Bacillus* spp. to produce pectinases along cocoa fermentation (H. G. Ouattara et al., 2008). Besides, the inoculated *Lp. plantarum* Lb2 and *Lm. fermentum* Lb1 presented a remarkable potential to metabolise carbohydrates as several glycoside hydrolases, glycosyl transferases and carbohydrate esterases were annotated in their genomes (Almeida et al., 2021). The participation of microorganisms in enzymatic activity in cocoa fermentation seems to be complementary to the endogenous hydrolases, mostly acting as activators of these effectors along fermentation as no enzymatic activity was directly and statistically correlated to the total microbiota.

A satisfactory cocoa fermentation is crucial for the development of aroma and taste of chocolate. For this, microbiota stimulates the biochemical transformations inside the beans, leading to the formation of desirable precursors (Castro-Alayo et al., 2019). A study reported the inoculation of defined starters elevated the levels of acids and esters in fermented beans, while decreased the amounts of aldehydes and ketones, and alcohols (Moreira et al., 2021). The opposite was observed in this study, the starters inoculation slightly augmented the levels of alcohols. Aldehydes and ketones were discreetly diminished in F0 (0h to 96h) and F1 (0h to 96h), and augmented in F2 (0h to 96h), while in F3 no changes were observed. Esters were also detected and were enhanced (0h to 96h) in F0, F1, F2, and no changes were observed in F3. Alcohols are not specifically related to quality of fermentation as they reflect the microbial metabolism along the process, while aldehydes and ketones play a role in the development of flavour due to their carbonylic compounds. Following, esters are the second most important compounds for cocoa fermentation and are related to the fruity-like aroma in fermented beans (Aprotositae et al., 2015).

The tendency of increasing desirable sensorial compounds (aldehydes, ketones, and esters) was observed at the end of all fermentations but was accompanied also by the emergence of detrimental metabolites: wine- and malty-like compounds. Besides, the expression of *luxS* of *Lp. plantarum* was associated with detrimental metabolites for fermented seeds quality. However, the absence of statistical significance of difference among the fermentations entangles the delimitation of the relationship of QS-related bacteria and the quality of cocoa fermentation.

### Quorum sensing and importance for fermentation

The importance of QS for bacterial stability and dominance in cocoa fermentation was first hypothesised by our research group. By the monitoring of a spontaneous cocoa fermentation carried out in Bahia state of Brazil, a metagenomic analysis predicted several taxa implied with interspecific QS, with evidence to lactobacilli (Almeida et al., 2020). In that work, it was proposed that novel studies could shed light into microbial ecology of cocoa fermentation by the evaluation of QS. As no *Acetobacter* spp. present *luxS* genes (Almeida et al., 2020; Almeida et al. 2022), the survey in this study was performed only for lactobacilli. It was observed *luxS* genes were highly expressed in F1 fermentation, most because of the inoculation of a high-density of lactobacilli cells. QS occurs in high-cell densities, and it is responsible for modulation of specific phenotypes that may confer adaptive advantages in harsh environments (Schluter et al., 2016). Initially, it was argued that QS could play a significant role in bacterial shifts in cocoa fermentation, impacting microbial succession (Almeida et al., 2020).

The data raised by this study brings a new vision, as it demonstrates the activity of QS along fermentation as argued previously (Almeida et al., 2020) and reveals the main player in this process: *Lp. plantarum*. While in F0 the bacterial diversity was low and in F2 and F3 fermentations there was a dominance of *Pantoea* genus, the addition of C2 cocktail induced the prevalence of lactobacilli, probably replacing *Enterobacteriaceae* members, which could confer stability and safety to the final product. It is worth highlighting that the *Pantoea* members were previously correlated with *luxS* genes (Almeida et al., 2020), and their replacement by *Lp. plantarum* could indicate a sum of forces that might proportionate *Lp. plantarum* dominance in detriment of *Enterobacteriaceae*.

By the other hand, in the view of some authors, the dominance of LAB in fermentation could be detrimental, due to the higher loads of lactic acid produced. Since it is a non-volatile compound, it may impact the taste of chocolate. Besides, research groups evidence the absence of LAB is not impeditive for fermentation (Ho et al., 2015, 2018). However, there is no consensus in literature about LAB detrimental properties, as many authors advocate these bacteria could be useful to replace undesirable microorganisms (Marwati et al., 2021) and provide good aroma notes (Viesser et al., 2020).

It’s worth to evidence of a pangenome survey based on 404 and 63 genomes of *Lp. plantarum* and *Lm. fermentum* species showed that *Lp. plantarum* species present multiple *luxS* gene homologues distributed in six gene clusters, while *Lm. fermentum* species present two gene clusters (Almeida et al., 2021). This can explain why *Lm. fermentum luxS* gene expression was incipient in comparison to *Lp. plantarum* genes. The augmented number of *luxS* genes in *Lp. plantarum* genomes could enhance its gene expression and confer adaptive fitness to outperform other lactobacilli in the same environment. Future studies focusing on primer design to monitor other lactobacilli species along cocoa fermentation are needed to compare the *luxS* expression patterns.

Still it is hard to affirm QS has an influence on cocoa fermentation quality as no significant changes were determined by the addition of QS-related bacteria in fermentation and changes in VOCs. Besides, no significant association of enzymatic activities and lactobacilli was found. As shown, QS could be related only to dominance of *Lp. plantarum*, while the quality is a sum of factors involving all microbiota in that environment and the intrinsic activity of cotyledons endogenous hydrolases (De Vuyst & Weckx, 2016). At the same time, more studies are needed to understand if LAB really impacts positively or negatively the fermentation, and if negatively, the disruption of QS communication could help to diminish lactobacilli loads in fermentation, especially of *Lp. plantarum* species. On the other hand, QS could aid the augmentation of lactobacilli to replace detrimental microorganisms.

## Conclusion

This was the first study to monitor enzymatic QS activities along cocoa fermentations. Although novel data was provided, this study was impacted by the intrinsic limitation of lab-scale fermentations, which not always can capture the real conditions of field such as the presence of a in house microbiota (from cocoa boxes, vessels, banana leaves and utensils), regional temperature and humidity and cocoa mass volume. Nevertheless, the findings were corroborated by literature in terms of microbial diversity and the microbial shifts during fermentation. The activity of *luxS* genes in all fermentations, but especially in that with an extra repertoire of lactobacilli, demonstrate the previous predictions of a metagenomic study, suggesting these experiments could be performed once in field conditions to compare the same observations. Moreover, it was impossible to fully link QS with cocoa fermentation quality, as additional tests are needed. However, the expression of *luxS* by lactobacilli *in situ* conditions is an interesting finding, as if future studies demonstrate LAB are detrimental to the process, due to lactic acid accumulation, strategies of QS disruption may be employed to limit these microorganisms. On the other hand, if these bacteria are crucial to the process, the stimulation of QS signalling would be useful to standardise the process favouring bacterial dominance. Finally, another limited knowledge in cocoa fermentation science is the absence of data regarding the microbial enzymatic activities and their extent in cocoa fermentation. This work showed and hypothesised that the main enzymatic activities related to changes in beans are displayed by endogenous hydrolases of the seeds, as no significant correlation of enzymes profiles and microbial composition was obtained. The data provided in this work could inspirate future studies to fill these new gaps.

## CRediT authorship contribution statement

**O.G.G. Almeida:** Conceptualization, Formal analysis, Investigation, Methodology, Writing original draft, Writing - review & editing**. M.G. Pereira:** Conceptualization, Methodology, Writing - review & editing**. R. L. Bighetti-Trevisan:** Methodology, Writing, - review & editing**. E. Santos:** Methodology. **E.G. De Campos:** Methodology. **G.E. Felis:** Methodology, Writing - review**. L.H.S. Guimarães:** Methodology, Writing - review. **M.L.T.M. Polizeli:** Methodology, Writing – review. **B.S. De Martinis:** Methodology, Writing – review. **E.C.P. De Martinis:** Conceptualization, Methodology, Writing – review & editing, Resources, Supervision.

## Conflict of interests

The authors declare no conflict of interests.

## Funding

This work was supported by a grant from The São Paulo Research Foundation, Brazil (FAPESP, grant #18/13564-3). O.G.G.A. is also grateful to FAPESP for Ph.D. fellowships (Processes # 17/13759-6 and # 18/26719-5). This research was also was financed in part by the Coordenação de Aperfeiçoamento de Pessoal de Nível Superior – Brasil (CAPES), Finance Code 001. ECPDM is grateful for a Researcher Fellowship from CNPq (PQ-2, Proc.# 306330/2019-9).

## Supplementary material

**Fig. A.1**. pH and Temperature monitoring measured in experimental fermentations.

**Fig. A.2.**
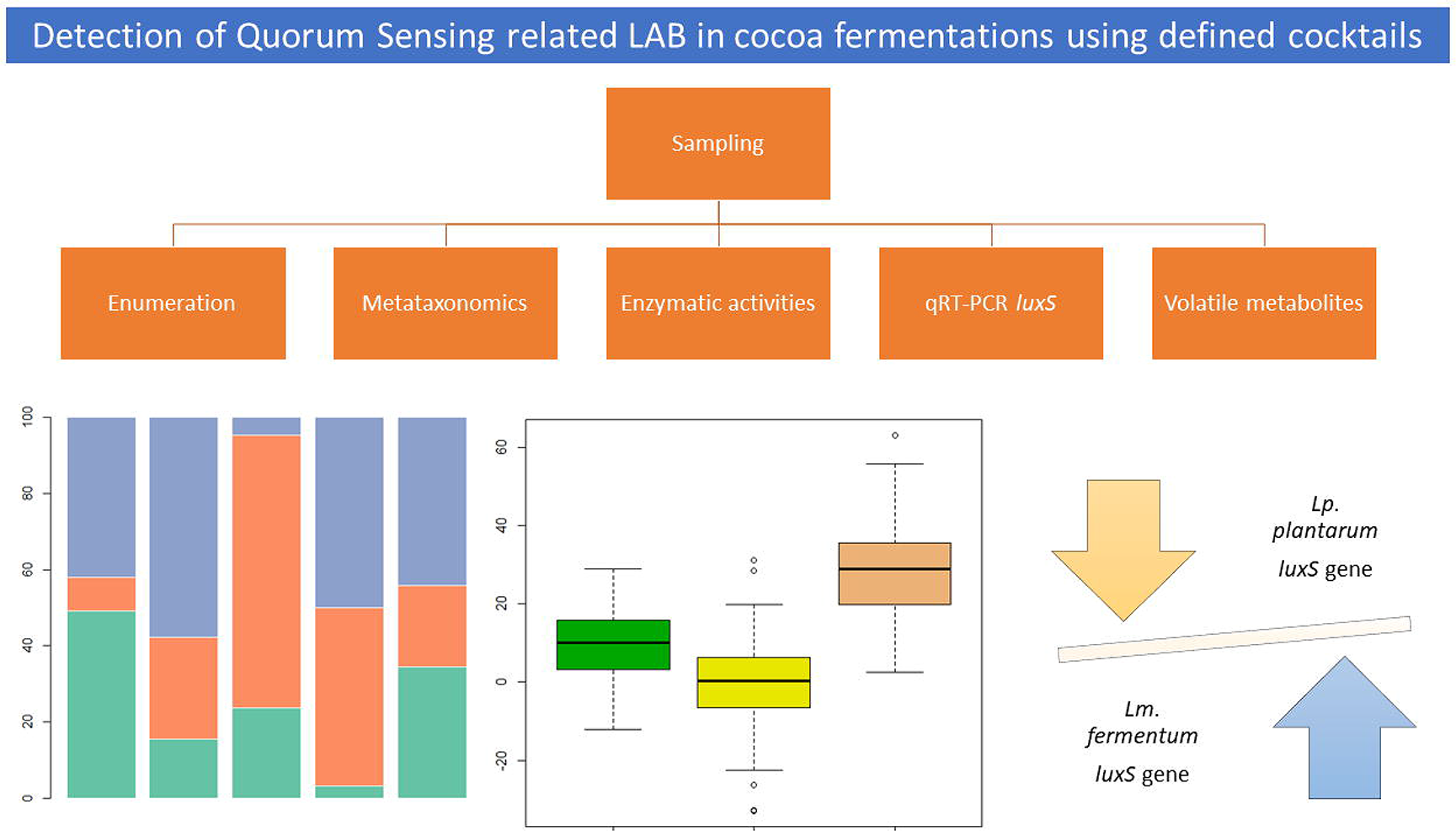
Kendall correlation among sensorial profiles and *luxS* gene expression of *Lp. plantarum* (luxS_Plan) and *Lm. fermentum* (luxS_Ferm) species.

## Notes

### Competing Interest Statement

The authors have declared no competing interest.

### Summary of Updates

Correction of abstract: Fermentations description.

