## Supplementary figures and images for "Investigating interspecific quorum sensing influence on cocoa fermentation quality through defined microbial cocktails"

### Supplementary Data

**Supplementary Fig. A.1.**

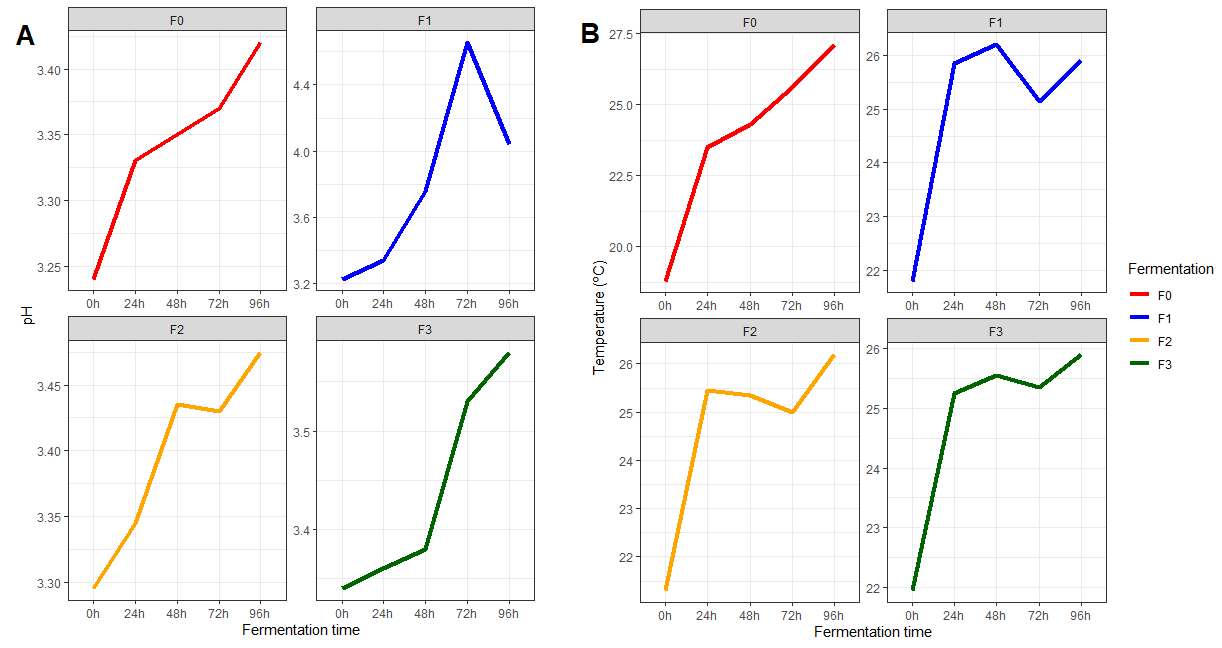

**Supplementary Fig. A.2.**

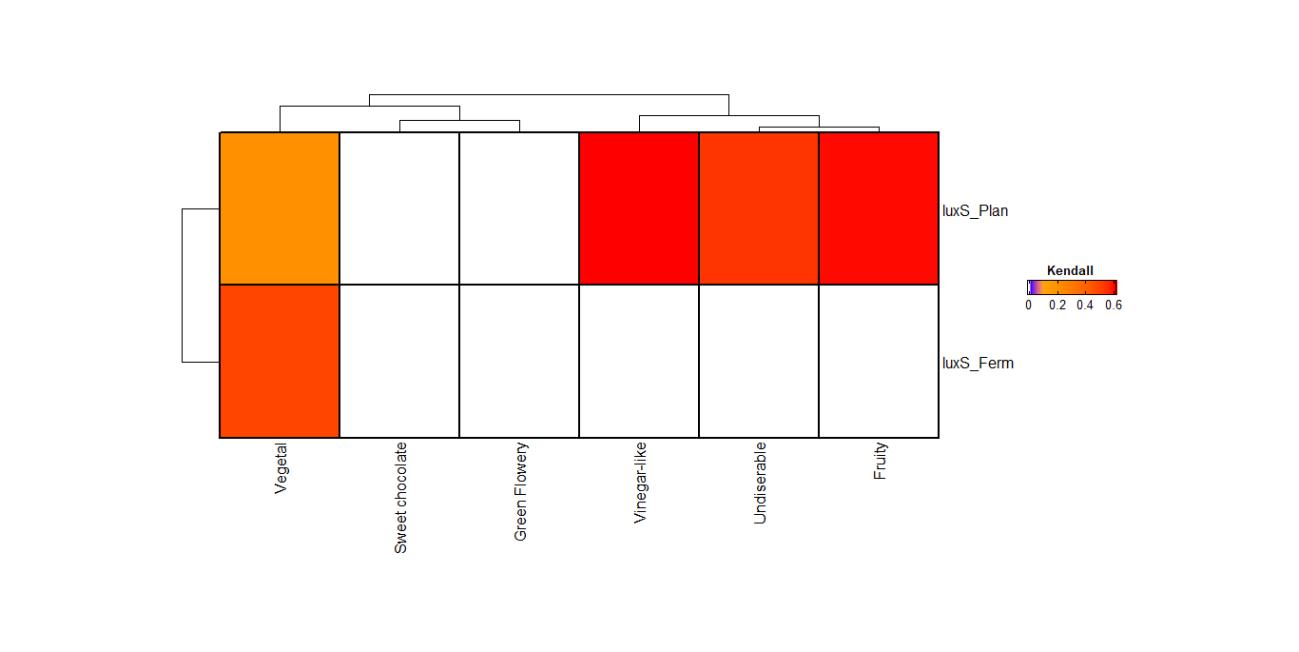
